# FAK-SFK Signaling Integrates ECM Rigidity Sensing and Engagement of ERBB2 to Activate YAP and Promote Invasive Growth and Metastasis in Breast Cancer

**DOI:** 10.1101/2022.08.03.502677

**Authors:** Xiaobo Wang, Shimin Wang

## Abstract

We have examined the mechanism through which fibrotic extracellular matrices promote tumor progression and metastasis in HER2+ breast cancer. We found that integrin-mediated mechano-transduction and engagement of ERBB2/ERBB3 cooperate to induce activation of YAP and invasive growth in stiff 3D Matrigel-Collagen I. Mechanistic studies revealed that joint activation of SRC Family Kinases (SFKs) by FAK and ERBB2/3 results in tyrosine phosphorylation and inactivation of LATS1/2 and MOB1. The ensuing activation of YAP enables HER2+ breast cancer cells to proliferate and invade in 3D Matrigel-Collagen I. In addition, tyrosine phosphorylation of LATS1/2 and MOB1 and activation of YAP are required for v-SRC-mediated transformation of fibroblasts. Finally, preclinical studies indicated that FAK-SRC signaling is required for primary tumor growth and lung metastasis in the MMTV-*Neu* mouse model of HER2+ breast cancer. Congruently, administration of dasatinib significantly increased the capacity of lapatinib to inhibit primary tumor growth and lung colonization in MMTV-*Ne*u mice. These findings indicate that integrin-mediated mechano-transduction functions as a rheostat to regulate ERBB2 signaling to YAP and suggest that co-targeting ERBB2 and SFKs may exhibit therapeutic efficacy in HER2+ breast cancer.

**Significance statement:** Stromal stiffness, which ensues from increased deposition and crosslinking of linear collagens, promotes oncogenesis and tumor progression in the breast. Moreover, tumor fibrosis is increased in the more aggressive subtypes of breast cancer, such as Triple Negative (TN) and HER2+ breast cancer. However, the mechanisms through which extracellular matrix stiffness regulates intracellular signaling are poorly understood. In this study, we show that the Focal Adhesion Kinase (FAK)-SRC Family Kinase (SFK) complex integrates the sensing of matrix rigidity by integrins and the activation of ERBB2/3 to promote activation of YAP and that YAP is required for breast cancer growth and invasion. Mechanistic studies demonstrate that SFKs can phosphorylate and inhibit LATS1 and MOB1, leading to activation of YAP. Simultaneous inhibition of ERBB2/3 with lapatinib and SFKs with dasatinib profoundly inhibits primary tumor growth and metastasis in a mouse model of HER2+ breast cancer. These findings suggest that the therapeutic efficacy of combinatorial blockade of ERBB2 and integrin-mediated mechano-transduction should be tested in biomarker-driven clinical trials.

## Introduction

Breast cancer has surpassed lung cancer and become the most common cancer type worldwide (1). Human Epidermal growth factor Receptor 2 (HER2)-positive breast cancer comprises 15-20% of all breast cancers (2). HER2 positivity confers more aggressive biological behavior and poorer clinical outcome (3). Although HER2-targeted therapy has been a favorable milestone in breast cancer treatment, durable drug responses are not consistently obtained due to the emergence of drug resistance (4). It is therefore important to discover novel mechanisms underlying tumor progression and metastasis and novel therapeutic vulnerabilities, to identify new biomarkers for therapy, and to develop rational combinations of targeted agents.

It has been shown that tumor fibrosis enhances tumor cell survival, growth, and migration, and facilitates the creation of a hypoxic and immune-suppressive tumor microenvironment (5). In breast cancer, increased deposition, linearization, and crosslinking of collagens renders the more aggressive Triple Negative (TN) and HER2+ subtypes of breast cancer stiffer than the less aggressive Luminal A and B subtypes (6). During breast cancer progression, increased extracellular matrix (ECM) stiffness promotes a proliferative and invasive phenotype in cancer cells, enabling them to invade through the basement membrane (BM) and dissemination to begin (7, 8). Tumor stiffness evaluated by breast elastography is found to be a predictive biomarker for poorer response to adjuvant chemotherapy in breast cancer. The high stiffness is significantly corelated with lower clinical complete response (9). In HER2 positive breast cancer, ECM rigidity drives resistance to HER2 inhibitor lapatinib (10). In Triple Negative breast cancer, chemotherapy induces Collagen IV upregulation in the ECM, which in turn leads to drug resistance and promotes cell invasion (11).

Integrin-mediated adhesion and signaling via focal adhesion kinase (FAK) and Src family kinases (SFKs) is critical for matrix stiffness-induced mechano-transduction, which relays and translates the mechanical cues from the matrix into biochemical signals, including the pro-mitogenic and pro-survival signaling pathways – Ras-ERK, PI3K/AKT, and YAP/TAZ pathways (12). The Hippo pathway effectors YES-associated protein (YAP) and transcriptional coactivator with PDZ-binding motif (TAZ, also known as WWTR1) are regulated by the mechanical cues and mediate cellular responses to ECM stiffness (13, 14). When MCF10A cells are shifted from stiff to soft matrix, YAP and TAZ translocate from the nucleus to the cytoplasm and become inactivated (14). The higher-grade tumor subtypes HER2+ and TN mammary tumors both show much higher nuclear levels of the YAP than the lower-grade Luminal A/B tumors, which is consistent with the stiffness levels (6). One of the key mechanisms of YAP/TAZ activation in response to mechanical signals is through Rho GTPase (Rho kinase, ROCK) that regulates the formation of actin bundles, stress fibers and contractile actomyosin structures via phosphorylating myosin light chain (MLC). Rho activity and cytoskeletal tension induced by stiff matrix are required for YAP/TAZ activation (15). However, the detailed biochemical and molecular mechanisms of how integrin-mediated mechano-transduction regulates Hippo signaling downstream effectors is still unclear.

The Hippo pathway, initially elucidated in Drosophila melanogaster, is a highly conserved signal transduction pathway that plays important roles in organ size control and tumor suppression (16-18). In certain conditions such as high cell density, extracellular matrix stiffness and lack of nutrients, the Hippo pathway is activated, with MST and LATS successively phosphorylated with the support of SAV1 and MOB1. Then, the activated LATS phosphorylates transcriptional co-activator YAP and TAZ, which prevent them from entering the nucleus to active gene expression (19-22). In transgenic mouse models, Loss of YAP suppresses oncogene-induced tumor growth (23). However, evidence surrounding the clinical implications of the Hippo pathway in breast cancer cases is scarce, and little is known about how the underlying molecular and biochemical mechanisms of this pathway are regulated.

Here we demonstrate that YAP tends to localize in the nucleus in high rigidity compared to soft matrix in HER2+ breast cancer cells while both HER2 inhibition and FAK/SFKs inhibition block YAP nucleus entering and suppress YAP activity. Moreover, we found that Src phosphorylates YAP upstream kinases LATS1 and MOB1, leading to inactivation of LATS1 and MOB1 and activation of YAP, which promotes cell invasive growth and tumor metastasis. SFK inhibitor dasatinib blocks this process and its combinational administration with lapatinib shows promising preclinical efficacy in the treatment of HER2+ breast cancer.

## Results

### ECM stiffness acts via HER2 to regulate FAK and YAP

YAP functions as an essential effector of mechano-transduction to regulate cell proliferation and differentiation (13, 24). GTPase RAP2 has been shown to mediate nuclear-cytoplasmic shuttling of YAP in response to ECM stiffness (14), but what upstream signals trigger the event remains unclear. The rigidity of HER2 positive breast tumors is higher than that of Luminal subtypes (6). To test if ECM stiffness acts through HER2 to regulate YAP, we compared the effects of ECM rigidity on YAP localization in HER2 positive breast cancer cell line SK-BR-3. In contrast to higher stiffness (40 kPa), lower stiffness (1 kPa) induced robust cytoplasmic translocation of YAP. We then activated HER2 receptor by adding different concentrations of the ligand Neuregulin. Immunofluorescence staining revealed that in soft matrices, where YAP remained in the cytoplasm, low doses of Neuregulin was enough to drive YAP from the cytoplasm into the nucleus (Fig. 1A). Immunoblotting also showed Neuregulin activated FAK, YAP and YAP downstream effectors, CYR61 and CTGF (Fig. 1B). To inactivate HER2, we used the HER2 inhibitor lapatinib. Lapatinib promoted cytoplasmic translocation of YAP in SK-BR-3 cells seeded in stiff matrices. Same results were also found in MMTV-Neu, a mouse Her2 positive cell line (Fig. 1C). Consistent with the staining data, immunoblotting showed Lapatinib dramatically reduced FAK and YAP activity and the expression of downstream targets (Fig. 1D). To study the clinical connection between HER2 and YAP, we examined patient dataset and found that high YAP mRNA level correlated with poor prognosis only in HER2 positive breast cancer compared to other subtypes (Fig. 1E and Fig. S1). These results indicate HER2 is an important mediator to activate FAK and YAP in response to tumor rigidity.

**Figure 1.**
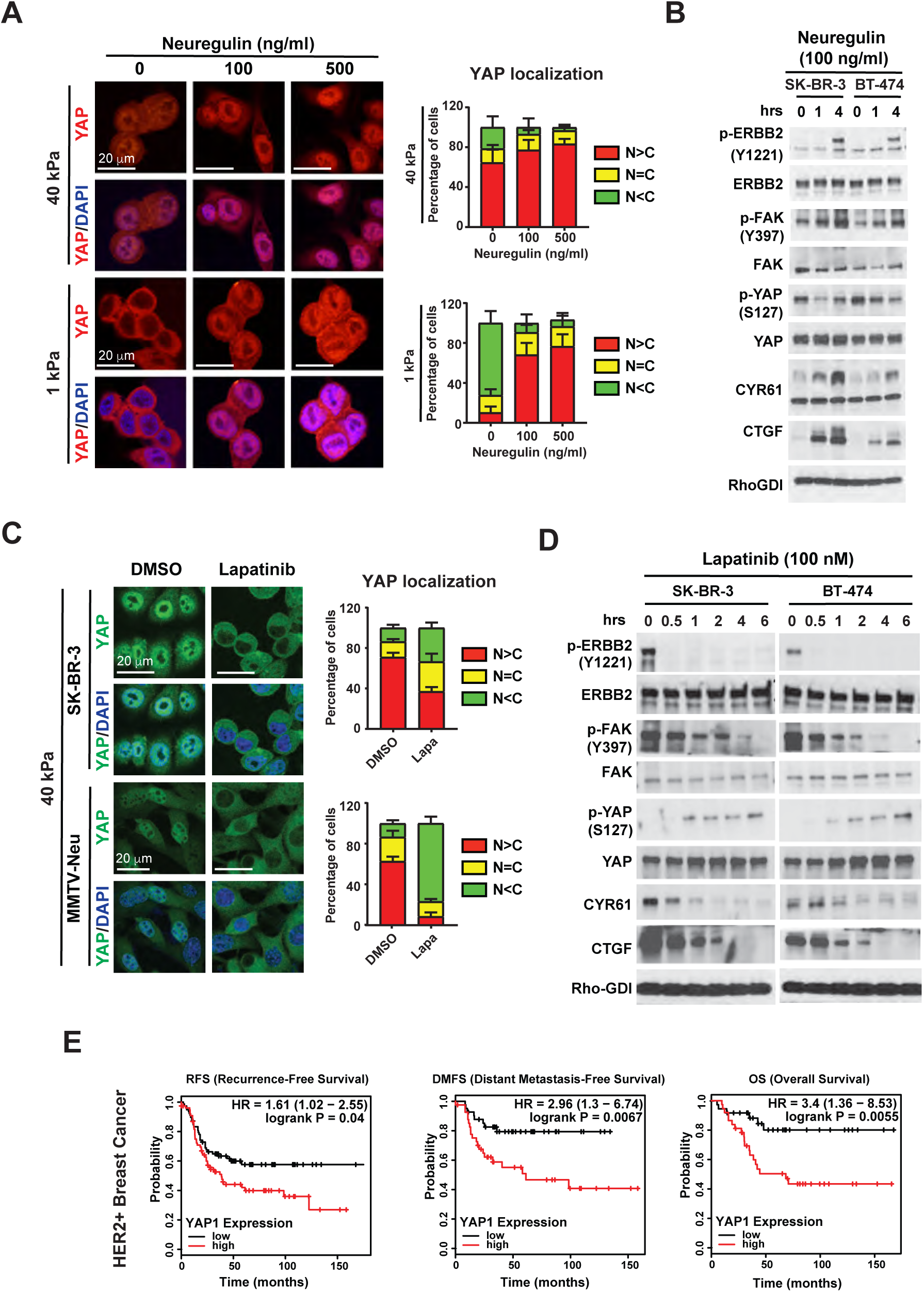
HER2 induces activation of FAK and YAP. (A) Immunofluorescence imaging and quantification of YAP localization in SK-BR-3 cells treated with different concentrations of Neuregulin in stiff (40 kPa) and soft (1 kPa) matrices. Error bars denote Mean + SD; Scale Bars, 20 µm. (N < C, less YAP in nucleus than in cytoplasm; N = C, similar levels of YAP in cytoplasm and nucleus; N > C, more YAP in nucleus than in cytoplasm.) (B) Immunoblotting with indicated antibodies in SK-BR-3 cells and BT-474 cells treated with Neuregulin (100 ng/ml) at different time points. (C) Immunofluorescence imaging and quantification of YAP localization in SK-BR-3 cells and MMTV-Neu cells treated with Lapatinib (100 nM) in stiff matrices (40 kPa). Error bars denote Mean + SD; Scale Bars, 20 µm. (D) Immunoblotting with indicated antibodies in SK-BR-3 cells and BT-474 cells treated with Lapatinib (100 nM) at different time points. (E) Kaplan-Meier plot showing the higher expression of YAP1 is correlated with reduced RFS (Recurrence-Free Survival), DMFS (Distant Metastasis-Free Survival), and OS (Overall Survival) in HER2-positive patients.

### Src induces tyrosine phosphorylation of LATS1

Immunoblotting data showed FAK activation can be induced by Neuregulin while inhibited by lapatinib (Fig. 1B and D). Previous studies have shown that HER2 activates FAK by forming a direct protein-protein interaction with the FAK-FERM-F1 lobe (25). Considering that FAK functions as a critical mediator of integrin signaling (12), we hypothesized that the integrin pathway acts through FAK to activate YAP in HER2 positive cells. To test it, adhesion assays were carried out in SK-BR-3 cells. Cells were suspended and then plated on Fibronectin for different time points. Immunoblotting results showed matrix adhesion induced FAK and YAP activation (Fig. 2A). These findings suggest the integrin signaling activates YAP in response to matrix adhesion. To examine the mechanism through which the integrin pathway regulates Hippo pathway, Co-IP experiment was performed to check the relation between the integrin signaling components and Hippo pathway members. Immunoblotting data showed FAK and Src, which belong to the integrin pathway, and LATS1, which is included in Hippo pathway, were in a same complex when plated on Fibronectin, whereas disassociated when treated with the FAK inhibitor PF271 (Fig. 2B). Since both FAK and Src are tyrosine kinases, we speculated if a direct phosphorylation existed. Surprisingly, we found Src, but not FAK, could phosphorylate LATS1 (Fig. S2A). Moreover, YAP was not phosphorylated by neither FAK nor Src directly (SI Appendix, Fig. S2B). Further analysis indicated that the constitutively active form of Src, Src-Y527F, induced a more robust signal than wild type Src, whereas the kinase dead form, Src-K295R, did not lead to LATS1 phosphorylation (Fig. 2C). Consistently, the Src inhibitor Saracatinib, but not FAK inhibitor PF271, could block Src-induced LATS1 phosphorylation (Fig. 2D). In vitro assay using purified GST-Src showed that the phosphorylation of LATS1 by Src was direct (Fig. S2C). To examine whether the phosphorylation was conserved, Src kinase family members Yes and Fyn, and LATS1 homolog protein LATS2 were tested. All Src family proteins phosphorylated both LATS1 and LATS2 (Fig. S2D). These findings confirm that Src induces tyrosine phosphorylation of LATS.

**Figure 2.**
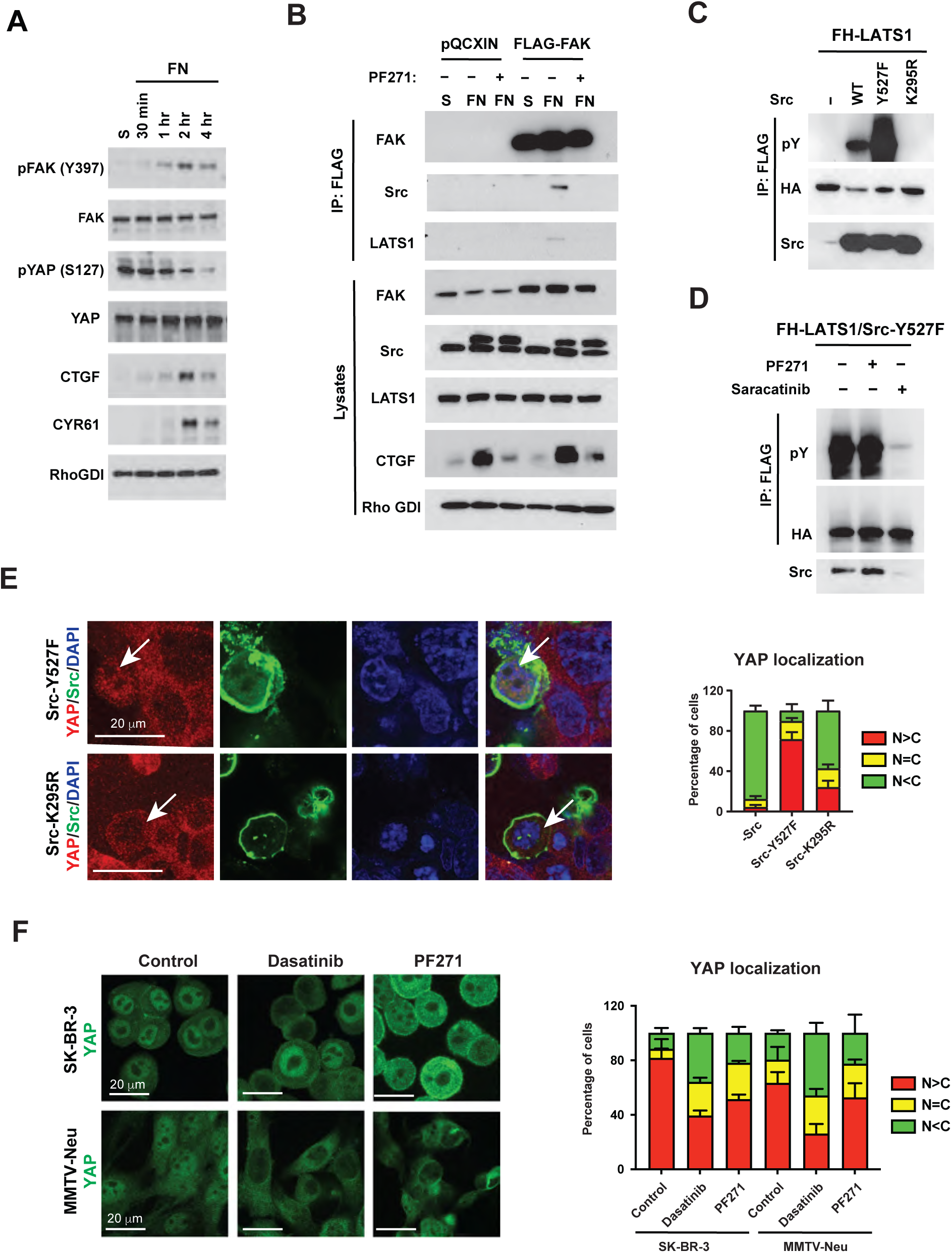
Src induces tyrosine phosphorylation of LATS1. (A) Immunoblotting with indicated antibodies in SK-BR-3 cells subjected to Fibronectin (FN) at different time points. (B) 293T cells were seeded in suspension or on Fibronectin with DMSO or PF271, and co-transfected with pQCXIN or pQCXIN-FLAG-FAK and Src-GFP. Immunoprecipitation was performed followed by immunoblotting with indicated antibodies. (C) FH-LATS1 and Control vector, wild-type Src, constitutively active form Src-527F, or kinase dead form Src-295R were co-expressed in 293T cells. FH-LATS1 was immunoprecipitated and pY was detected by immunoblotting. (D) FH-LATS1 and Src-527F were co-expressed in 293T cells. After the treatment with FAK inhibitor PF271 or Src inhibitor Saracatinib, FH-LATS1 was immunoprecipitated and pY was detected by immunoblotting. (E) Immunofluorescence imaging and quantification of YAP localization after expressing Src-Y527F or Src-K295R in 293A cells. Scale Bars, 20 µm. (F) Immunofluorescence imaging and quantification of YAP localization after treatment with Dasatinib or PF271 in SK-BR-3 and MMTV-Neu cells. Scale Bars, 20 µm.

To test whether LATS1 phosphorylation by Src can change YAP localization, we performed the immunofluorescence staining in 293A cells and found that co-expression of active Src-Y527F, but not kinase dead Src-K295R, dramatically increased YAP nuclear translocation, suggesting Src promotes YAP activity in a kinase-dependent manner (Fig. 2E). To test whether FAK and SFKs are required in the YAP activation induced by integrin-mediated mechanotransduction in HER2 breast cancer, we treated SK-BR-3 and MMTV-Neu cells with dasatinib and PF271 in stiff matrices. Both dasatinib and PF271 induced YAP translocation into the cytoplasm, but dasatinib had more robust effects (Fig. 2F). Taken together, these results demonstrate that integrin signaling activates YAP via FAK/SFKs, and Src-induced LATS1 phosphorylation is a crucial mechanism for YAP activation.

### Mapping phosphorylation sites on LATS1

To determine the phosphorylation sites on LATS1, two rounds of Mass Spectrometry analysis were carried out and 23 potential phosphorylated sites were found in total (Table S1 and Fig. 3A). According to the ratio of mod/unmod and the conservation, Y916, Y1026 and Y1076 were chosen to be tested. First, we introduced single site mutations by mutating the tyrosine to phospho-dead phenylalanine (Y916F, Y1026F, Y1076F) and examined the effetcts. When co-transfected with Src-Y527F, each single mutation form of LATS1 showed decreased tyrosine phosphorylation level, indicating each site contributes to the phosphorylation. Next, combinational mutation on all the three sites (3YF) was tested and the tyrosine phosphorylation was completely blocked (Fig. 3B). These results demonstrate that mutation of all these three sites prevent Src-induced LATS1 phosphorylation. To further understand the roles of these phosphorylation sites, the antibodies specifically targeting the phosphorylated LATS1 on each site were made and examined. Dot blotting showed all three antibodies worked well (Fig. S3A). Consistently, the phosphorylation specific antibodies showed Src phosphorylated all three sites on wild type LATS1, but not mutant LATS1-3YF (Fig. 3C). After the validation, we tested the functions of LATS1 phosphorylation at these three sites. LATS1/2 were knocked out in 293A cells, and wild type and different mutation forms of LATS1 were re-introduced. YAP tended to localize in the nucleus when LATS1/2 were knocked out. Re-introduction of wild type LATS1 was effective to drive YAP out of the nucleus. However, re-introduction of LATS1-3YE, which mutated three Y sites to E, to mimic the highly-phosphorylation state, could not induce the cytoplasmic translocation of YAP (Fig. 3D). In comparison, the re-introduction of single mutation versions, LATS1-Y916E, -Y1026E, and -Y1076E, only partially changed the YAP localization (Fig. S3B-C). To further confirm this conclusion, we preformed the same assays in MMTV-Neu cells and got the similar results (Fig. S3D). These findings suggest that the Y916/Y1026/ Y1076 together contribute to LATS1 phosphorylation and its inhibitory effects on YAP.

**Figure 3.**
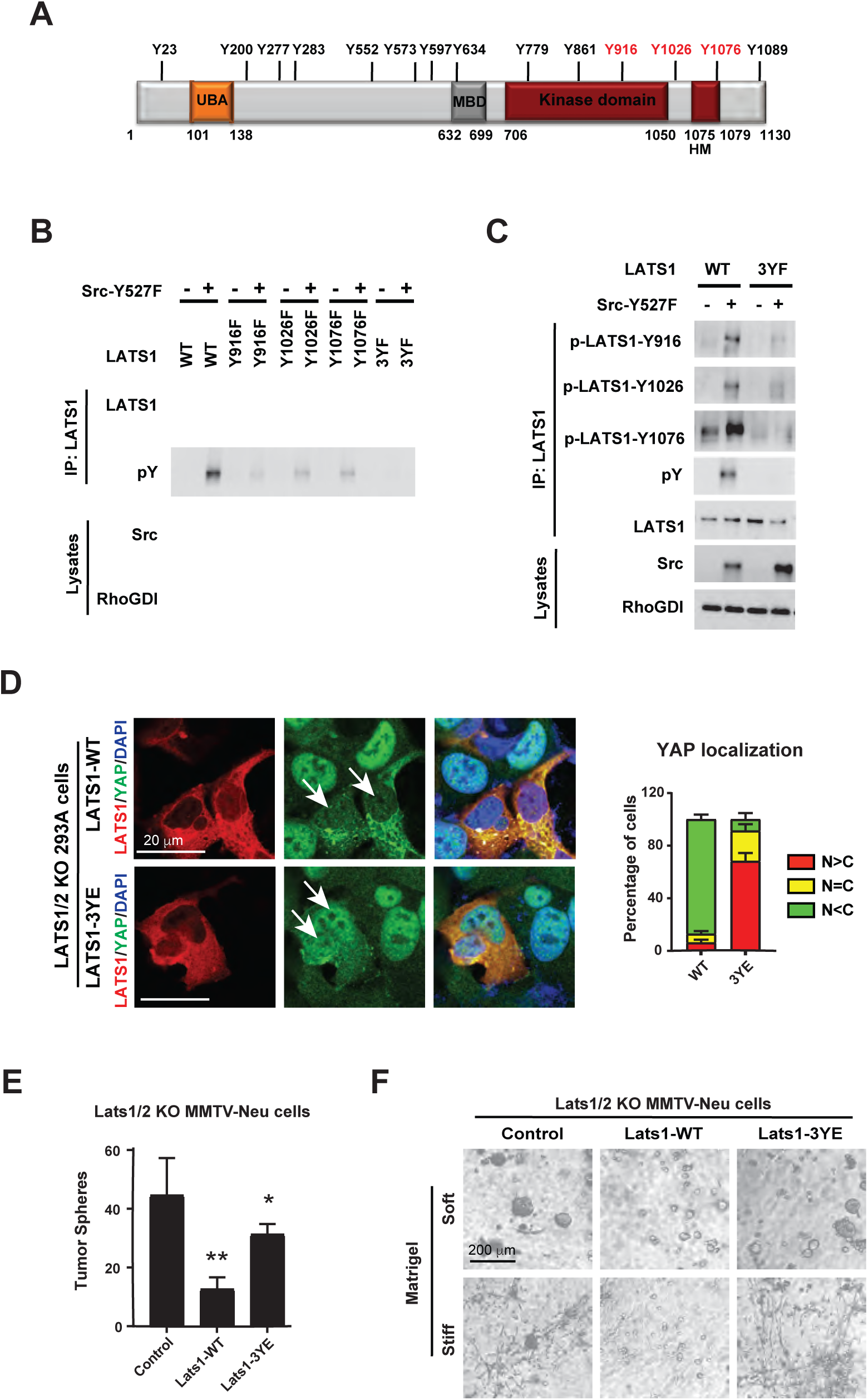
Src induces LATS1 phosphorylation at Y916/Y1026/Y1076. (A) A schematic illustration of domains and motifs of LATS1 and Tyr phosphorylation sites identified in Mass spectrometry mapping. (B) 293T cells were co-transfected with Src-527F and LATS1 (wild type and different mutant forms, 3YF: Y916/1026/2076F), then LATS1 was immunoprecipitated followed by immunoblotting with indicated antibodies. (C) 293T cells were co-transfected with Src-527F and LATS1 (wild type or 3YF), then LATS1 was immunoprecipitated followed by immunoblotting with phosphorylation site specific antibodies. (D) Immunofluorescence imaging of HA-LATS1 (RFP) and YAP (GFP) and quantification of YAP localization in LATS1/2 knockout 293A cells re-introduced with wild type LATS1 or mutant LATS1-3YE (Y916/1026/2076E). Scale Bars, 20 µm. (E) Quantification of tumor sphere assay of Lats1/2 knockout MMTV-Neu cells re-introduced with Lats1-WT or Lats1-3YE. Error bars denote Mean + SD; P values (two-tailed t-test): *p<0.05, **p<0.01. (F) Invasive cell growth assay of Lats1/2 knockout MMTV-Neu cells re-introduced with Lats1-WT or Lats1-3YE on Soft Matrigel (Matrigel only) and Stiff Matrigel (Matrigel with collagen). Scale Bars, 200 µm.

To further corroborate the roles of LATS1 phosphorylation in biological functions of tumor cells, tumor sphere and invasive growth assays on Matrigel with or without collagen were performed. When wild type and mutant Lats1 was introduced in Lats1/2 knockout MMTV-Neu cells, tumor sphere formation was suppressed by wild type Lats 1 but not mutant Lats1-3YE (Fig. 3E). Similarly, on soft and stiff Matrigel, only wild type Lats1 but not Lats1-3YE suppressed clone size and branching respectively (Fig. 3F). These results suggest the conclusion that Src-induced tyrosine phosphorylation of LATS1 at Y916/Y1026/Y1076 has great impacts on YAP-mediated cell growth, invasion, and cell stemness traits.

### Src induces tyrosine phosphorylation of MOB1

FAK has been shown to directly phosphorylate MOB1, a scaffold protein as part of the complex with LATS1 in the Hippo core kinase cascade, to activate YAP (26). To examine if this also applies to our system, MOB1 were co-transfected with activated Src or FAK in 293T cells, and tyrosine phosphorylation levels were tested. To our surprise, no obvious phosphorylation signal was detected when MOB1 was co-expressed with FAK, but a robust phosphorylation band appeared in the co-expression with Src, indicating Src also phosphorylated MOB1 (Fig. 4A). To test if Src or FAK could phosphorylate other Hippo pathway components, MST1/2 and SAV were co-expressed with Src or FAK, but no phosphorylation was observed (Fig. S4A). The previous studies demonstrated MOB1-Y26 was phosphorylated by FAK (26) and p-MOB1-Y26 antibody was commercially available. Therefore, MOB1-Y26F mutation was made and tested first. Constitutively active Src, but not kinase dead Src, phosphorylated both wild type MOB1 and mutant MOB1-Y26F, suggesting Y26 was not the right site or not the only site for phosphorylation by Src (Fig. 4B). Given MOB1 is a relatively small protein with only has 8 tyrosine sites, we performed single point mutagenesis for all the 8 sites individually and tested one by one. In the immunoblotting, none of the single mutations blocked MOB1 phosphorylation by Src, suggesting multiple tyrosine phosphorylation sites existed on MOB1 (Fig. S4B). To identify the exact sites, all 8 tyrosine sites were mutated to phenylalanine, which was MOB1-8YF, and then each phenylalanine was mutated back to tyrosine one by one and tested again. MOB1-8YF largely blocked the phosphorylation signal, but when mutated back, each mutation showed contributions to the phosphorylation, indicating activated Src phosphorylated all tyrosine sites on MOB1 (Fig. S4C). To study the functions of MOB1 phosphorylation on YAP, wild type MOB1 and mutant MOB1-Y8E were re-introduced in MOB1A/B knockout 293A cells. Re-introduction of wild type MOB1 was effective to drive YAP out of the nucleus. However, re-introduction of MOB1-8YE could not induce the cytoplasmic translocation of YAP (Fig. 4C). Then we evaluated the roles of MOB1 phosphorylation in cell growth and invasion in Mob1 re-introduced Mob1/2 knockout MMTV-Neu cells. On soft and stiff Matrigel, only wild type Mob1 but not Mob1-8YE suppressed cell growth and invasion (Fig. 4D). These results suggest that MOB1 phosphorylation induced by Src has great impacts on YAP-mediated cell growth and invasion. Although no direct phosphorylation of LATS1/MOB1 by FAK were found, whether FAK directly affects Src-induced phosphorylation was tested. FAK or siFAK were co-expressed with Src in 293T cells, tyrosine phosphorylation levels of LATS1 and MOB1 were detected. No obvious changes of the phosphorylation levels were found, suggesting FAK had no direct influences on Src to phosphorylate LATS1 and MOB1 (Fig. S4D).

**Figure 4.**
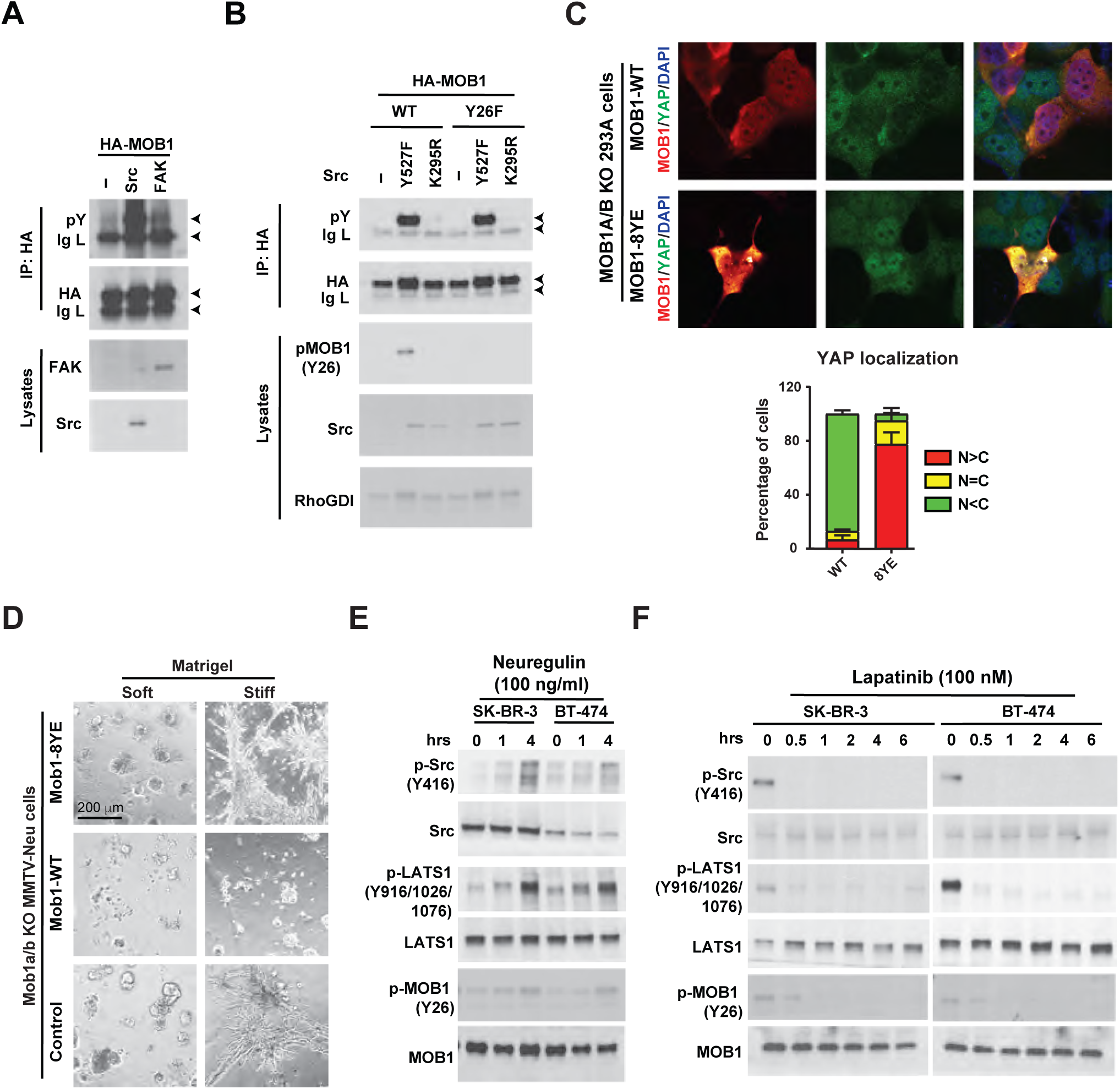
Src induces tyrosine phosphorylation of MOB1. (A) 293T cells were transfected with HA-MOB1 together with Src or FAK. HA-MOB1 was Immunoprecipitated followed by immunoblotting with indicated antibodies. (B) Wild type HA-MOB1 or mutant HA-MOB1-Y26F was co-expressed with Src in 293T cells, followed by immunoprecipitation and immunoblotting with indicated antibodies. (C) Immunofluorescence imaging of YAP (GFP) and HA-MOB1 (RFP) and quantification of YAP localization in MOB1A/B knockout 293A cells re-introduced with wild type MOB1 or mutant MOB1-8YE (8YE: All eight tyrosine sites of MOB1 mutated to glutamic acid). Scale Bars, 20 µm. (D) Invasive cell growth assay of Mob1a/b knockout MMTV-Neu cells re-introduced with wild type Mob1 or Mob1-8YE on Soft and Stiff Matrigel. Scale Bars, 200 µm. (E) Immunoblotting with indicated antibodies in SK-BR-3 cells and BT-474 cells treated with Neuregulin at different time points. (F) Immunoblotting with indicated antibodies in SK-BR-3 cells and BT-474 cells treated with Lapatinib at different time points.

To examine whether HER2 activation affects Src activity and LATS1/MOB1 tyrosine phosphorylation levels, Neuregulin was added into SK-BR-3 cells and BT-474 cells at different time points. Upon HER2 activation, Src was phosphorylated at Y416, and tyrosine phosphorylation levels of LATS1 and MOB1 increased (Fig. 4E). Consistently, when treated with HER2 inhibitor lapatinib, Src activity was suppressed, and phosphorylation levels of LATS1 and MOB1 decreased (Fig. 4F). These results suggest that HER2 regulates Src activity to phosphorylate LATS1 and MOB1. Taken together, these findings demonstrate that in HER2 positive breast cancer, HER2 activates Src, thus phosphorylating Hippo pathway components LATS1 and MOB1, and activating YAP.

### B77 requires Src-induced LATS1/MOB1 phosphorylation to activate YAP

Since Src has been identified to be the kinase that induces phosphorylation of LATS1 and MOB1, we further tested our findings in the v-Src-transformed B77 cells. Both datatinib and PF271 could inhibit the nuclear translocation of YAP in stiff matrices (Fig. 5A). The RNA-seq analysis showed that YAP signature was enriched in DMSO-treated B77 cells compared to datatinib-treated cells (Fig. S5A). These results suggest that Src activity is required for YAP activation in B77 cells. Consistent with what we found in breast cancer cell lines, only wild type Lats1 but not phosphorylation-mimetic Lats1-3YE could inhibit YAP activation (Fig. 5B) and suppress tumor sphere formation and invasive growth (Fig. 5C-D) in Lats1/2 knockout B77 cells. Similarly, only wildtype Mob1 but not phosphorylation-mimetic Mob1-8YE inhibited the tumor invasive growth in both soft and stiff matrices (Fig. 5E). To test the functions of YAP in B77 cells, we knocked down YAP and performed MTT assays and soft agar assays. YAP knockdown significantly inhibited cell proliferation (Fig. S5B) and colony formation (Fig. 5F). Moreover, the constitutively active form YAP-5SA largely increases the sensitivity to Src inhibitor Saracatinib in B77 cells (Fig. 5G). These results demonstrate YAP activation is crucial for the Src-induced transformation.

**Figure 5.**
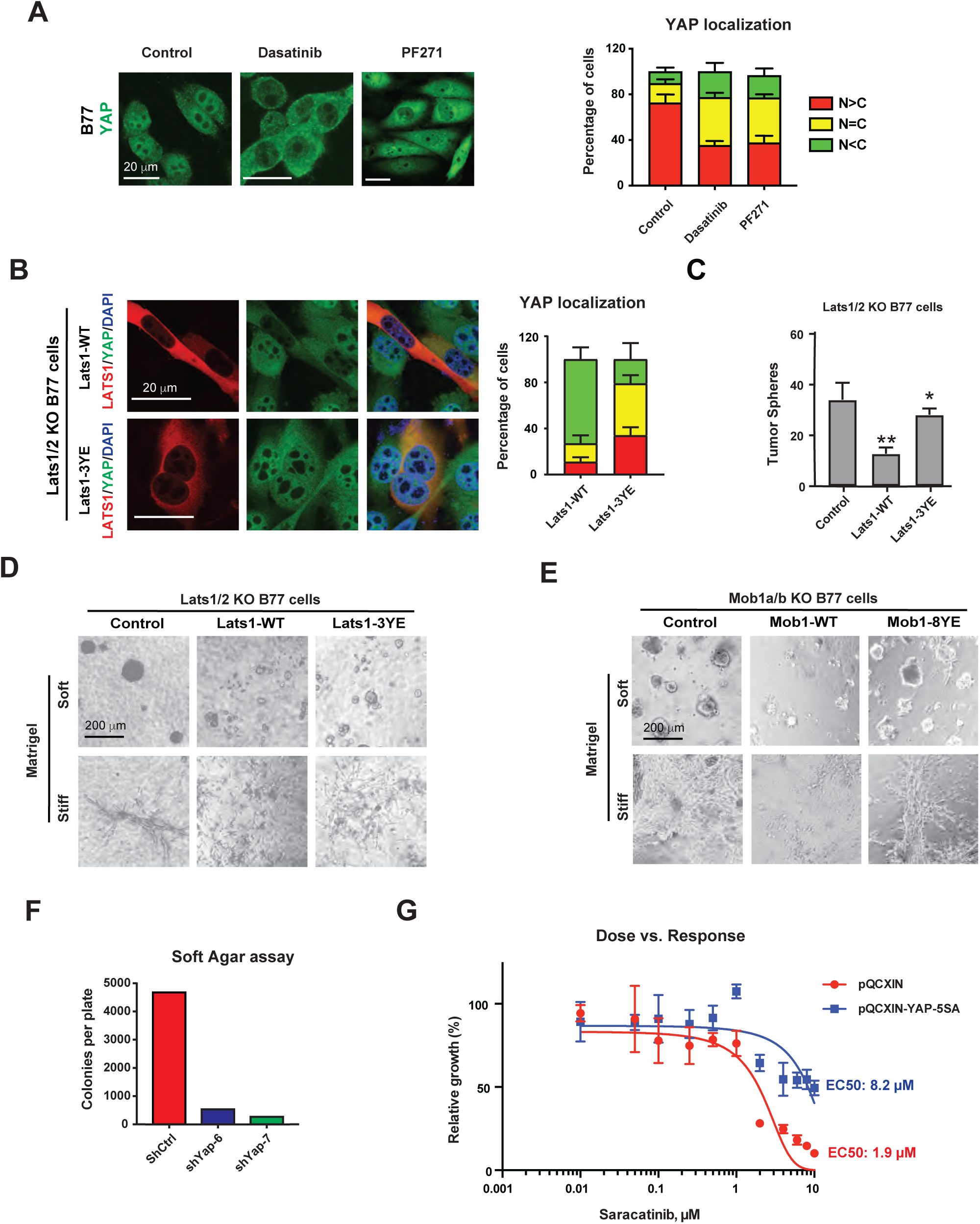
YAP activation is required for Src-induced transformation. (A) Immunofluorescence imaging and quantification of YAP localization after treatment with Dasatinib or PF271 in B77 cells. Scale Bars, 20 µm. (B) Immunofluorescence imaging and of YAP (GFP) and HA-LATS1 (RFP) and quantification of YAP localization in LATS1/2-knock out B77 cells re-introduced with LATS1-WT and LATS1-3YE. Error bars denote Mean + SD; Scale Bars, 20 µm. (C) Quantification of tumor sphere assay in Lats1/2 knockout B77 cells re-introduced with Lats1-WT and Lats1-3YE. Error bars denote Mean ± SD; P values (two-tailed t-test): *p<0.05, **p<0.01. (D) Invasive cell growth assay of Lats1/2 knockout B77 cells re-introduced with Lats1-WT and Lats1-3YE on Soft and Stiff Matrigel. Scale Bars, 200 µm. (E) Invasive cell growth assay of Mob1a/b knockout B77 cells re-introduced with wild type Mob1 or Mob1-8YE on Soft and Stiff Matrigel. Scale Bars, 200 µm. (F) Soft agar assays in Yap knockdown B77 cells. (G) Dasatinib sensitivity test in B77 cells transfected with pQCXIN or pQCXIN-YAP-5SA.

### Depletion of Yap/Taz and Fak inhibits cancer growth and metastasis

To explore the biological effects of Yap/Taz depletion in breast cancer cells, both in vitro and in vivo functional assays were performed. Yap/Taz knockdown was carried out by doxycycline inducible shRNA and the efficiency was evaluated by immunoblotting (Fig. S6A). Firstly, to test the cell growth and proliferation rate, MTT assays were conducted in MMTV-Neu cells. Individual knockdown of Yap and Taz suppressed cell growth, whereas the proliferation rate was further reduced when Yap and Taz were knocked down together (Fig. 6A). Consistently, in soft agar assays, knockdown of Yap/Taz suppressed the colony formation more robustly than individual knockdown of Yap or Taz in MMTV-Neu cells (Fig. 6B). These results indicate Yap and Taz have redundant functions. Next, shYap/Taz MMTV-Neu cells were placed on soft Matrigel (Matrigel only) and stiff Matrigel (Matrigel with collagen) for growth. Robust reductions in colony formation and branching growth were observed under soft and stiff conditions respectively after Yap/Taz knockdown (Fig. 6C). To examine the roles of Yap and Taz in cell stemness, tumor sphere assays were conducted in MMTV-Neu cells. Knockdown of Yap/Taz dramatically decreased tumor sphere number and size (Fig.6D). These results suggest that Yap and Taz depletion inhibits cell growth, invasion, and stemness traits in Her2+ breast cancer cells. To study the function of Yap and Taz in tumor growth and metastasis *in vivo*, inducible knockdown of Yap and Taz in MMTV-Neu-TGL cells was generated and injected into mammary fat pad or tail vein of female nude mice. In orthotopic injection experiments, after two weeks of growth, when the tumor volume reached approximately 100 mm^3^, doxycycline was added to induce the knockdown of Yap and Taz. Tumor growth was monitored. After two weeks of doxycycline treatment, mice were euthanized, and tumors were collected. In doxycycline treated group, the tumor growth was significantly inhibited (Fig. 6E). The findings indicate that Yap/Taz depletion suppresses primary tumor growth of Her2+ breast cancer. In the metastatic studies, doxycycline was added to induce knockdown of Yap and Taz one week after the tail vein injection. Bioluminescence signals in lungs were significantly weaker in Yap/Taz knockdown group (Fig. 6F). The results demonstrate that depletion of Yap/Taz suppresses metastatic outgrowth in lung of Her2+ breast cancer.

**Figure 6.**
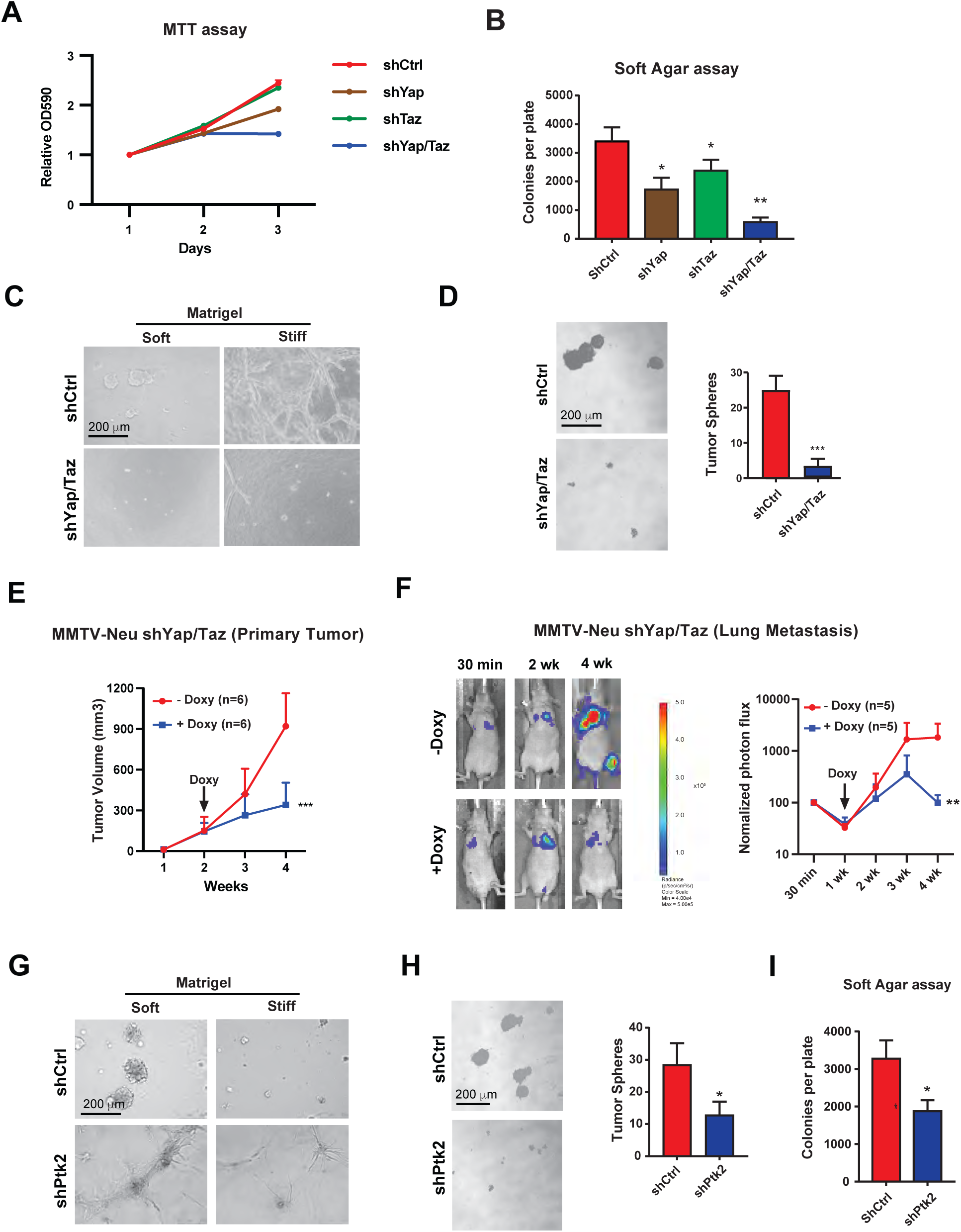
Depletion of Yap/Taz and Fak inhibit tumor growth and invasion in vitro and in vivo. (A) MTT assays in Yap/Taz knockdown MMTV-Neu cells. (B) Soft agar assays in Yap/Taz knockdown MMTV-Neu cells. Error bars denote Mean ± SD; P values (two-tailed t-test): *p<0.05, **p<0.01. (C) Invasive cell growth assay on Soft Matrigel (Matrigel only) and Stiff Matrigel (Matrigel with collagen) in Yap/Taz knockdown MMTV-Neu cells. Scale Bars, 200 µm. (D) Tumor sphere assay and quantification in Yap/Taz knockdown MMTV-Neu cells. Error bars denote Mean + SD; P values (two-tailed t-test): ***p<0.0001. Scale bars, 200 µm. (E) Quantification of tumor volumes of female nude mice injected in mammary fat pad with inducible Yap/Taz knockdown MMTV-Neu cells. Doxycycline induction began at Week 2 after cell injection. Error bars denote Mean + SD; P values (two-tailed t-test): ***p<0.0001. (F) Representative Images and normalized photon flux of nude mice injected through tail vein with inducible Yap/Taz knockdown MMTV-Neu cells. Doxycycline induction began at Week 1 after cell injection. Error bars denote Mean + SD; P values (two-tailed t-test): **p<0.01. (G) Invasive cell growth assay on Soft Matrigel (Matrigel only) and Stiff Matrigel (Matrigel with collagen) in Ptk2 knockdown MMTV-Neu cells. Scale Bars, 200 µm. (H) Tumor sphere assay and quantification in Ptk2 knockdown MMTV-Neu cells. Error bars denote Mean + SD; P values (two-tailed t-test): *p<0.05. Scale bars, 200 µm. (I) Soft agar assay quantification in Ptk2 knockdown MMTV-Neu cells. Error bars denote Mean + SD; P values (two-tailed t-test): *p<0.05.

Although no direct phosphorylation of LATS1 and MOB1 induced by FAK was found, pharmacological inhibition of FAK could inhibit YAP activation (Fig. 2F). Therefore, we evaluated the effects of genetic perturbation of FAK in breast cancer. We knocked down Ptk2, the gene that encodes Fak, in MMTV-Neu cells. Knockdown efficiency after doxycycline induction was confirmed by immunoblotting (Fig. S6A). Functional cell assays were performed to determine the effects of Fak depletion. In soft and stiff matrices, the cell growth in Matrigel was suppressed by Fak knockdown (Fig. 6G). The colony numbers on soft agar also decreased after Fak knockdown (Fig. 6H). Consistently, knockdown of Fak led to the reduction in size and number of tumor spheres (Fig. 6I). To test the effects of Fak knockdown on primary tumor growth and metastasis *in vivo*, MMTV-Neu-TGL cells expressing inducible shPtk2 were injected in female nude mice. Fak knockdown slowed tumor growth in primary mammary tumors (Fig. S6B) and lung metastases (Fig. S6C). These results support the conclusion that Fak is important for cell growth, invasion, and metastasis in Her2 positive breast cancer. To further explore the connection between FAK and Hippo signaling from these *in vivo* studies, samples from metastatic tumors were collected for RNA extraction followed by RNA-seq analysis. The KEGG pathway analysis revealed that lots of known pathways associated with FAK signaling were enriched, such as tight junction, focal adhesion, and cell adhesion molecules. In addition, Hippo signaling pathways were also enriched, suggesting that Fak functions upstream of Hippo signaling to regulate tumor growth and metastasis (Fig. S6D).

### Dasatinib shows preclinical efficacy in the treatment of Her2 positive breast cancer and metastasis

To test preclinical efficacy of inhibition of FAK/SFK signaling to YAP in HER2 positive breast cancer, we used the Her+ breast cancer mouse model MMTV-Neu, which began to develop spontaneous breast tumors 3 months after birth. When tumors reached 2 mm diameter, mice were treated with multi-kinase inhibitor dasatinib and FAK inhibitor PF271 respectively (Fig. 7A). 24 hours after the treatment, a group of mice was euthanized, and tumors were collected for target inhibition tests. IHC staining for Ki67 and C-Caspase3 in the harvested tumors showed decreased cell proliferation and increased cell apoptosis after the treatment with dasatinib and PF271 (Fig. S7A-B). Hippo signaling markers Yap and p-Yap were also examined. Yap was mainly in the nucleus in control group while Yap tended to localize in the cytoplasm in dasatinib and PF271 treated groups. Consistently, p-Yap increased after the treatment with dasatinib and PF217, which indicated the suppressive activity of Yap (Fig. S7C). Tumor sizes were monitored until the euthanasia after 3 weeks of treatment. Both dasatinib and PF271 slowed down the tumor growth (Fig. 7B). Based on the model that YAP is activated by both matrix adhesion and activated HER2 in MMTV-Neu mice, we tested the clinical grade SFK inhibitor dasatinib in combination with HER2 inhibitor lapatinib. We anticipated cooperation between the 2 drugs in each combination even if HER2 is the sole activator of FAK/SRC in these tumors, as hitting 2 steps in the same linear cascade has proven efficacious in the clinic (27). For primary tumors, MMTV-Neu-TGL cells were injected into mammary fat pad, and treated with vehicle, lapatinib, dasatinib, or lapatinib combined with dasatinib. Drug treatments began once tumors reached approximately 100 mm^3^. Tumor volumes were monitored during the treatment. Single treatment with lapatinib or dasatinib inhibited tumor growth, while lapatinib and dasatinib combination group showed slowest tumor growth rate (Fig. 7C). For metastatic tumors, MMTV-Neu-TGL cells were injected through tail vein, and treatments began after one week. Bioluminescent imaging and statistical analysis revealed that both lapatinib and dasatinb alone suppressed growth in lung metastases, but combination of the two drugs further improved the effects (Fig. 7D). Next, the lung metastases samples were collected and subjected to IHC staining. Compared to control group, Ki67 decreased and C-Caspase3 increased in all treatment groups, especially in the combinational group with lapatinib and dasatinib (Fig. 7E, upper and Fig. S7D). Moreover, Yap and p-Yap were examined. It was clearly that Yap localized in the nucleus in control group whereas Yap was mainly in the cytoplasm in lapatinib and dasatinib combinational treatment group. Yap activity decreased as p-Yap increased in the combinational treatment group (Fig. 7E, lower). Based on these results, we concluded that SFK inhibitor dasatinib in combination with lapatinib shows promising preclinical efficacy to treat HER2 positive breast cancer.

**Figure 7.**
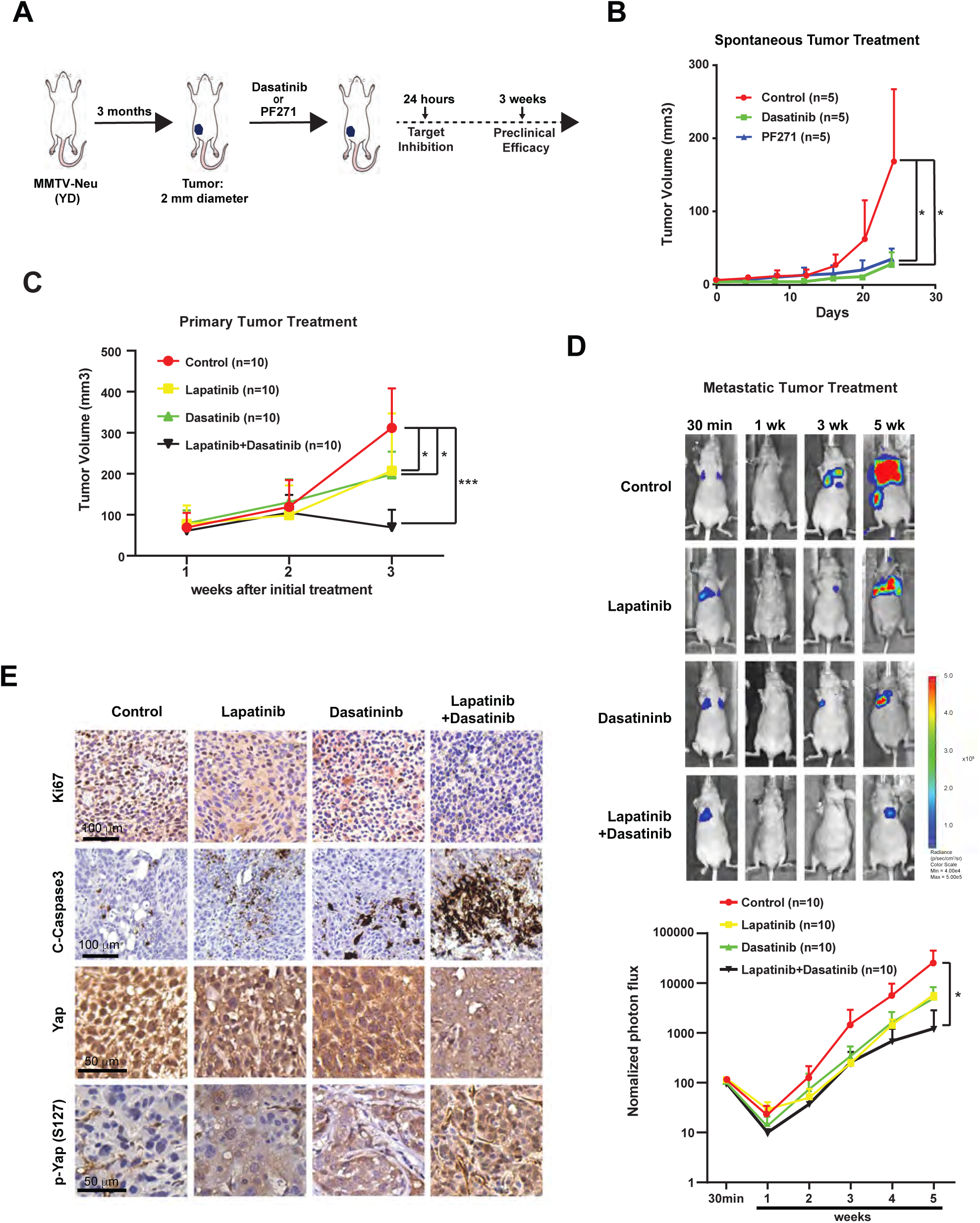
Dasatinib inhibits Her2 breast cancer growth and metastasis. (A) A schematic illustration of workflow of inhibitor treatment in MMTV-Neu mice model. (B) Quantification of tumor volume of MMTV-Neu mice treated with vehicle, Dastinib (20 mg/kg, oral gavage, qd) or PF271 (50 mg/kg, oral gavage, qd). Error bars denote Mean + SD; P values (two-tailed t-test): *p<0.05. (C) Quantification of tumor volumes of female nude mice injected in mammary fat pad with 300,000 MMTV-Neu-TGL cells after the treatment with vehicle, Lapatinib (50 mg/kg, oral gavage, qd), Dasatinib (50 mg/kg, oral gavage, qd), and Lapatinib with Dasatinib. Treatment began once tumors reached approximately 100 mm^3^. Error bars denote Mean + SD; P values (two-tailed t-test): *p<0.05, ***p<0.0001. (D) Representative Images and normalized photon flux of nude mice injected through tail vein with 300,000 MMTV-Neu-TGL cells after the treatment with vehicle, Lapatinib (50 mg/kg, oral gavage, qd), Dasatinib (50 mg/kg, oral gavage, qd), and Lapatinib with Dasatinib. Treatment began 1 Week after the injection. Error bars denote Mean + SD; P values (two-tailed t-test): *p<0.05. (E) Representative of IHC staining imaging of Ki67, C-Caspase3, Yap, and p-Yap (S127) in the lung metastases samples harvested from MMTV-Neu cell-injected nude mice after the treatment indicated in Fig. 7D. Scale Bars as indicated.

## Discussion

Tumor stiffening occurs with the tumor pathogenesis and progression. The matrix is stiffer in HER2 positive and triple negative breast cancer subtypes than the less aggressive luminal counterparts (6). The mechanical inputs from the ECM are sensed by mechano-sensors at the cell membrane and relayed by mechano-transducers to promote YAP/TAZ activity at sites where mechanical forces are high (13). Consistently, we found that YAP is activated by rigid matrices HER2 positive breast cancer cells. To understand the mechano-transducers of YAP activation, we used pharmacological inhibition and genetic perturbation of the integrin signaling components and identified a critical role for integrin signaling via FAK-SRC in activation of YAP/TAZ in HER2 positive and triple negative breast cancer cells. In HER2 positive cells, integrin signaling synergizes with HER2 to activate YAP. This provides new evidence for the joint regulation between integrin signaling and RTKs. But what triggers the FAK/SFK and YAP activation in triple negative cancer is poorly known. Studies have shown the effects of integrin-FAK signaling on tumor initiation, metastasis, and resistance to chemotherapy in triple negative breast cancer. EGFR and integrin β4 cooperate to active FAK and trigger the tumor initiation and metastasis in triple negative breast cancer (28). A recent study in triple negative MDA-MB-231 cells shows that EZH2-upregulated integrin β1 activates FAK, which phosphorylates TGFβ receptor type I (TGFβRI) at Y182 to increases its binding to TGFβ receptor type II (TGFβRII), thus activating TGFβ signaling and bone metastatic potential (29). ECM deposition and remodeling promotes chemoresistance by inducing FAK activation through integrin α1/α2/α5 (11, 30). These studies shed light on FAK activation by upstream integrin and RTKs. However, whether YAP is activated and whether YAP activation is required for these biological functions in breast cancer still await further investigation.

Although ECM stiffening and YAP/TAZ activation in breast cancer have been clearly implicated in previous research and further validated in our studies, the biochemical mechanisms that link ECM-responsive integrin-FAK/SFK signaling and YAP/TAZ activation are largely unknown. We conducted co-immunoprecipitation and identified the interaction between FAK/Src and YAP upstream regulator LAST1. Following studies show Src can induce phosphorylation of LATS. Mass spectrometry and mutagenesis analysis revealed that Src phosphorylates LATS at 3 conserved sites (Y916, Y1026, and Y1076), leading to loss of the kinase activity of LATS1. To determine whether Src-induced LATS1 is the driver mechanism, CRISPR-Cas9-mediated knockout of LATS1/2 was followed by reconstitution with wild type or an unphosphorylatable (Y916/1026/1076F or 3YF) or a phosphorylation-mimetic (Y916/1026/1076E or 3YE) mutant of LATS. Only 3YE mutant but not wild type or 3YF can induce the YAP activation and malignancy of HER2 positive breast cancer cells. Another Hippo signaling component MOB1, can also be phosphorylated by Src. Contrary to the previous study, which demonstrates FAK phosphorylates MOB1, we did not see the FAK-mediated MOB1 phosphorylation (26). MOB1 is phosphorylated at 8 tyrosine sites and the phosphorylation-mimetic mutant MOB1-8YE can rescue the YAP activation and malignancy of MOB1A/B knockout HER2 positive and triple negative breast cancer cells. Both LATS and MOB1 phosphorylation by Src disable their kinase activity for YAP inhibition, thereby activating YAP. These results establish a novel link between FAK/SFK and YAP activation, in which LATS1and MOB1 are both phosphorylated by Src. In the Hippo signaling, LATS1 and MOB1 phosphorylation are mediated by MST. In addition to MST, LATS can be phosphorylated by MAP4Ks, PKA, CHK1/2, CDK1/CDC2, and Aurora A kinase (31). Opposed to that, MST-independent MOB1 phosphorylation is scarcely reported. EGFR promotes the phosphorylation of MOB1 and the inactivation of LATS1/2 independently of MST1/2. (32). Therefore, the identified Src-mediated direct phosphorylation on LATS and MOB1 expand the understanding of Hippo signaling.

We then used an inducible knockdown system to evaluate the biological functions YAP, TAZ, and FAK in HER2 positive breast cancer cells. FAK knockdown decreases cell proliferation and invasive growth and inhibits mammary tumor growth and metastasis in mice. YAP and TAZ function largely in a redundant fashion to sustain tumor growth and metastasis. The YAP/TAZ knockdown shows higher inhibitory effects than FAK knockdown. These results make it rational to test the effects of FAK and/or YAP/TAZ inhibition in breast cancer.

First, we tested the individual drug SFK inhibitor Dasatinib and FAK inhibitor PF271 in spontaneous MMTV-Neu mammary tumor mouse model. Both drugs suppressed YAP activation and slowed down the tumor growth. Then, for larger scale experiments, we injected MMTV-Neu-TGL cells into mammary fat pat and tail vein for the drug evaluation in primary and metastatic tumors respectively. Integrin signaling contributes to the resistance of HER2 positive SK-BR-3 and BT-474 cells plated on laminin-5 to HER2 inhibition. FAK inhibitor TAE226 can sensitize the cells to HER2 antagonists, Trastuzumab and Lapatinib (33). Therefore, the simultaneous inhibition of integrin-FAK/SFK and HER2 may improve the efficacy. In both primary and metastatic tumors, the single drug administration with dasatinib and lapatinib showed efficacious tumor-inhibitory effects. The combination of dasatinib and lapatinib was most effective. These preclinical studies demonstrate the synergistic anti-tumor effect between the blockade of integrin-FAK/Src-YAP axis and HER2. In addition, several studies have shown basal-like triple negative breast cancer are most sensitive to dasatinib compared to luminal subtypes (34-36). It will be important to evaluate the preclinical efficacy of FAK and SFK inhibitors alone or in combination with chemotherapy in triple negative breast cancer.

Taken together, this study demonstrates integrin-FAK/SFK mediates the mechano-transduction from stiff ECM and activates YAP, which is required for cell invasive growth and tumor metastasis; and unmasks the mechanisms through which YAP is activated by integrin signaling - Src induces phosphorylation of LATS and MOB1, leading to their inactivation and ensuing YAP activation. Furthermore, Integrin signaling and HER2 jointly regulate YAP activation and combinational inhibition of these two pathways shows promising preclinical efficacy in HER2-positive breast cancer.

## Acknowledgements

We thank Dr. Filippo Giancotti for conceiving the study, designing and analyzing the experiments, and providing suggestions on the manuscript writing. We thank Drs. Young-Mi Kim and Yan Jiang for performing and analyzing experiments. We thank Drs. Hong Chen and Hyunho Han for analyzing patient dataset and RNA-seq data.

## Author Contributions

X.W. and S.W. designed, performed, and analyzed the experiments, and wrote the manuscript under the supervision of Dr. Filippo Giancotti.

## EXPERIMENTAL MODEL AND SUBJECT DETAILS

### Cell Lines

SK-BR-3 and BT-474 were obtained from ATCC. SK-BR-3 and BT-474 cells were cultured in RPMI 1640 medium (Gibco) supplemented with Fetal Bovine Serum (FBS, 10%; Gibco) and Penicillin/Streptomycin (100 IU/ml). 293FT packaging cells were from Invitrogen and cultured according to manufacturer’s instructions. 293T, 293A and B77 cells were cultured in DMEM (Gibco) supplemented with FBS (10%) and Penicillin/Streptomycin (100 IU/ml). The mouse MMTV-Neu cell line was established in our lab as previously described (37). Briefly, MMTV-Neu cells were isolated from MMTV-Neu (YD) mice and dissociated by incubation with dispase (2.5 mg/ml) and collagenase type I (Invitrogen) for 2-4 hours and were then plated on Collagen I (20 µg/ml) in DMEM supplemented with FBS (10%) and Penicillin/Streptomycin (100 IU/ml). However, after three to four passages, these isolated cells lost expression of Neu (YD), presumably because of inactivation of the MMTV LTR promoter. To overcome this limitation, the cells were transduced with pWZL-Neu8142, which encodes oncogenic rat ErbB2. The transduced cell line was cultured in DMEM supplemented with FBS (10%) and Penicillin/Streptomycin (100 IU/ml). All cell lines were grown at 37 °C, 5% CO_2_, and 95% humidity.

### Compounds

The inhibitors purchased from Selleckchem were pre-prepared as a 10 mM stock solution in DMSO and used in *in vitro* cell assays. The inhibitors from MedChemExpress (MCE) were prepared in accordance with the manufacturer’s guidelines and used in *in vivo* mouse experiments. The drug concentration was adjusted in alignment with the dosage in mouse treatment. For Dasatinib, Defactinib, and Lapatinib, the following solvents were added to the drug individually and in order: 5% DMSO + 40% PEG 300 + 5% Tween 80 + 50% Saline. The final concentration of Dasatinib was 5 mg/ml or 12.5 mg/ml, Defactinib 25 mg/ml, and Lapatinib 12.5 mg/ml. For PF-562271 (PF271), the following solvents were added to the drug individually and in order: 10% DMSO + 40% PEG 300 + 5% Tween 80 + 45% Saline. The final concentration was 12.5 mg/ml.

### Mice

For all the animal studies in the present study, the study protocols were approved by the Institutional Animal Care and Use Committee (IACUC) of UT MD Anderson Cancer Center. Female nude mice (aged 6-8 weeks) were purchased from Jackson Laboratory. Female MMTV-Neu mice were generated as previously described (37). For primary tumor growth assay, cells were re-suspended in 100 µl of PBS with Matrigel in 1:1 ratio and injected into the fat pad of mammary gland number 3. The volume of the tumors was calculated as V = L x W x W/2, where L and W stand for tumor length and width, respectively. Experimental metastasis assays were performed as previously described (38, 39). Cells were re-suspended in 100 µl of PBS and injected into tail vein with a 26G tuberculin syringe. Metastatic burden was detected through non-invasive bioluminescence imaging of experimental animals using an IVIS Spectrum. To investigate the effects of drug treatment, compounds were delivered once a day through oral gavage, control group was treated with vehicle. Bioluminescence signal was measured weekly using the ROI tool in Living Image software (Xenogen) to verify successful injection and to monitor metastatic outgrowth.

### Inducible knockdown of gene expression

Inducible shRNAs constructs were generated Gene Editing & Screening Core (GES Core), Memorial Sloan Kettering Cancer Center. shRNAs were designed using the SplashRNA algorithm (40). A previously described optimized lentiviral miR-E expression backbone system was used for Dox inducible LT3RENIR (Neomycin, dsRed), LT3REPIR (Puromycin, dsRed), and LT3GEPIR (Puromycin, Green) expression (41).

### CRISPR/Cas9-mediated gene knockout

LentiCRISPRv2 vectors were used for the gene knockout (42, 43). Guide RNA (gRNA) oligos were designed on the CHOPCHOP website (http://chopchop.cbu.uib.no/) (44). The target site sequence was flanked on the 3’ end by an NGG PAM sequence. NGG PAM sequence was not included in the designed oligos. Each pair of oligos were phosphorylated and annealed, followed by the ligation by Quick Ligase (NEB) to BsmBI-digested and purified LentiCRISPRv2 (antibiotic markers: Puromycin or Blasticidin) backbone vectors.

### Tumor Sphere Assay

Single cell suspensions of cells (1,000 cells/ml) were plated on ultra-low attachment plates and cultured in serum-free PrEGM supplemented with B27 (1:50), basic fibroblast growth factor (bFGF, 20 ng/ml) and epidermal growth factor (EGF, 40 ng/ml) for 10 days. Tumor spheres were visualized under phase contrast microscope, photographed, and counted. For serial passage, tumor spheres were collected using 70-mm cell strainers and dissociated with Accutase at 37 °C for 30 minutes to obtain single cell suspensions.

### Matrigel 3D Culture

Dissociated cells were incubated in PrEGM medium supplemented with B27 (1:50), basic fibroblast growth factor (bFGF, 20 ng/ml) and epidermal growth factor (EGF, 40 ng/ml). Matrigel bed was made in 6-well plate by putting 4 separate drops of Matrigel per well (50 µl of Matrigel per drop). Plates were placed in 37 °C CO_2_ incubator for 30 minutes to allow the Matrigel to solidify. For each sample, 100 µl of cell suspension was mixed with 100 µl of cold Matrigel and pipetted on top of the Matrigel bed (50 µl each). The plates were then incubated at 37 °C for another 30 min. Warm PrEGM (2.5 ml) was then added to each well. The cells were cultured and monitored for 10-14 days with 50% medium change every 3 days. For immunostaining experiments, the cells were cultured in 8-well chamber slide. Cells were fixed with 4% paraformaldehyde for 20 minutes and proceeded to standard immunostaining protocol.

### Cell culture with polyacrylamide-based hydrogels

Glass coverslips were methacrylated in methacrylation buffer (20 ml of 100% Ethanol; 600 µl of 1:10 Acetic acid; 100 µl of Methacrylate) for 3 minutes. Slides were treated with DCDMS for 1 minute and removed with a KimWipe. Polyacrylamide gel solution was prepared according to the stiffness and polymerization, human placenta fibronectin (10 µg/ml) or rat tail collagen I (25–50 µg/ml) were used to coat the sulfo-sanpah-activated hydrogels according to the preferences of the cell lines. Hydrogels were rinsed with PBS 2 times and pre-incubated with culture medium for equilibration. Cells were seeded at 10,000 cells per well for high stiffness, and 20,000 cells per well for low stiffness in a volume of 680 µl per well (14).

### Immunoprecipitation and Immunoblotting

293T cells in 6-well plates were transiently transfected with 1 µg of pRK5 plasmid expressing FLAG-HA tagged LATS1 and 1 µg of plasmid expressing Src-Y527F by using Lipofectamine 3000 (Invitrogen). After 24 hours, the cells were lysed with RIPA buffer or RIPA buffer without SDS as indicated. To isolate FLAG-HA-LATS1, extracts were pre-cleared with agarose-mouse IgGs and incubated with anti-FLAG M2 Affinity Gel (Sigma) at 4 °C for 2 hours. The immunoprecipitates were washed four times with RIPA buffer or RIPA buffer without SDS as indicated, and bound proteins were dissociated in 20 µL of 1 x SDS sample buffer (25 mM Tris, pH 6.8; 4% SDS; 5% Glycerol; bromophenol blue). Eluted proteins were separated on 4-12% Bis-Tris SDS-PAGE gels (Invitrogen) and transferred to Immobilon-P membranes (Millipore). Membranes were incubated in blocking buffer (5% skim milk, 0.1% Tween, 10 mM Tris at pH 7.6, 100 mM NaCl) for 1 hour at room temperature and then with primary antibodies diluted in blocking buffer for another hour at the same temperature. After three washes, the membranes were incubated with goat anti-rabbit HRP-conjugated antibody or goat anti-mouse HRP-conjugated antibody at room temperature for 1 hour and subjected to chemiluminescence using ECL (Pierce #1856136). When indicated, cells were lysed in SDS boiling buffer (10 mM Tris, pH 7.5; 1% SDS; 50 mM NaF; 1 mM NaVO4) and subjected to SDS-PAGE gel electrophoresis and immunoblotting. Other immunoprecipitation experiments were carried out with the same protocols.

### Mass Spectrometry

Phosphorylated LATS was resolved using SDS polyacrylamide gel electrophoresis, followed by staining with Simply Blue and excision of the separated protein bands. *In situ* trypsin digestion of polypeptides in each gel slice was performed as described (45). All electrospray ionization experiments were performed using a QSTAR XL hybrid mass spectrometer (AB/MDS Sciex) hyphenated with microscale capillary reversed-phase HPLC [Famos autosampler (LC Packings), Agilent 1100 HPLC pump (Agilent)]. The columns were packed in-house using Magic C18 (5 µm, 200 Å, Michrom BioResources) beads. The buffer compositions are as follows: buffer A: 2.5% acetonitrile, 0.2% formic acid; buffer B: 2.5% H_2_O, 0.2% formic acid. For the quantitative experiments a 5-min gradient was used with mass spectra being acquired every 0.15 sec. Data analysis and quantitation was done using the Analyst software package provided by Applied Biosystems/MDS Sciex.

### Immunohistochemistry Staining

Immunohistochemistry on paraffin-embedded sections was performed using a Discovery XT processor (Ventana Medical Systems). The tissue sections were deparaffinized with EZPrep buffer, antigen retrieval was performed with CC1 buffer. Sections were blocked for 30 minutes with Background Buster solution (Innovex), followed by avidin-biotin blocking for 8 minutes (Ventana Medical Systems). Sections were incubated with anti-YAP (Invitrogen, #PA1-46189); anti-p-Yap (S127) (Invitrogen, #MA5-33207); anti-Ki67 (Abcam, # ab16667); anti-Cleaved Caspase 3 (Cell Signaling, # 9661) for 5 hours, followed by 60-minutes incubation with biotinylated horse anti-rabbit (Vector Labs, # PK6101) at 1:200 dilution. The detection was performed with DAB detection kit (Ventana Medical Systems) according to the manufacturer’s instruction. Slides were counterstained with hematoxylin and mounted using Permount (Fisher Scientific).

### Immunofluorescence staining

For immunofluorescence, cells were fixed in 4% formaldehyde/PBS for 10 minutes and then were treated with 0.1% Triton X-100 for 15 minutes at room temperature. After blocking, the cells were stained with anti -Yap (Sigma, #WH0010413M1); anti-Yap/Taz (Cell signaling, #8418) for 1 hour at room temperature, followed by 1-hour incubation with 488/561 conjugated goat anti-rabbit/mouse (Cell signaling, # 4412, #4413) at 1:200 dilution. Then cover samples with ProLong® Gold Antifade Reagent with DAPI (#8961) and coverslip. Allow slides to cure in the dark at room temperature for 24 hours, then seal the coverslips with nail polish. Images were captured with a Leica SP8 X Confocal Microscope and then were exported from Leica imaging software.

### RNA-seq Analysis

Total RNA was isolated using the TRIzol (Thermo Fisher Scientific) or RNeasy Mini kit coupled with RNase-free DNase set (Qiagen). Three replicates for each sample were generated and analyzed. Data were sequenced and analyzed by BGI. Libraries were prepared by using the standard methodology from Illumina and run on a HiSeq2500 system. Raw reads were quality-checked and subsequently mapped to the human or mouse genome using Tophat2 (2.2.4) with default settings (46). Differential gene expression was analyzed using the DESeq2 (1.8.1) package in R using default settings (47). Gene set enrichment analysis (GSEA) (48) was performed on a pre-ranked gene list that was generated based on the gene expression changes between the two groups. The hallmark gene sets, GO gene sets and KEGG pathways from the Molecular Signatures Database (MSigDB v5.1) (48) were evaluated by GSEA with 1,000 permutations, and those significantly (FDR < 0.1) enriched pathways and GO gene sets were reported using ggplot2 R package. Heatmap analysis was performed to show the gene expression patterns between groups, using heatmap3 R package with ward2 as distance function. Gene expressions in the heatmap were transformed in logarithm scale and normalized accordingly.

### Statistical analysis

All the statistical details including the statistical tests used, exact number of animals, definition of center, and dispersion and precision measures for the experiments can be found in the figures, figure legends or in the results. Statistical analyses were performed by using GraphPad Prism 8 software, with a minimum of three biologically independent samples for significance. For tumor injection experiments, each mouse was counted as a biologically independent sample. Results are reported as mean ± SD. Comparisons between two groups were performed by using an unpaired two-sided Student’s t-test (P < 0.05 was considered significant, * P<0.05, ** P<0.01, ***P<0.001). All experiments were reproduced at least three times, unless otherwise indicated.

## References

1. H. Sung et al., Global Cancer Statistics 2020: GLOBOCAN Estimates of Incidence and Mortality Worldwide for 36 Cancers in 185 Countries. CA Cancer J Clin 71, 209–249 (2021).

2. N. Cancer Genome Atlas, Comprehensive molecular portraits of human breast tumours. Nature 490, 61–70 (2012).

3. Y. Kun et al., Classifying the estrogen receptor status of breast cancers by expression profiles reveals a poor prognosis subpopulation exhibiting high expression of the ERBB2 receptor. Hum Mol Genet 12, 3245–3258 (2003).

4. S. Loibl, L. Gianni, HER2-positive breast cancer. Lancet 389, 2415–2429 (2017).

5. B. Piersma, M. K. Hayward, V. M. Weaver, Fibrosis and cancer: A strained relationship. Biochim Biophys Acta Rev Cancer 1873, 188356 (2020).

6. I. Acerbi et al., Human breast cancer invasion and aggression correlates with ECM stiffening and immune cell infiltration. Integr Biol (Camb) 7, 1120–1134 (2015).

7. O. Chaudhuri et al., Extracellular matrix stiffness and composition jointly regulate the induction of malignant phenotypes in mammary epithelium. Nat Mater 13, 970–978 (2014).

8. K. R. Levental et al., Matrix crosslinking forces tumor progression by enhancing integrin signaling. Cell 139, 891–906 (2009).

9. M. Hayashi et al., Evaluation of tumor stiffness by elastography is predictive for pathologic complete response to neoadjuvant chemotherapy in patients with breast cancer. Ann Surg Oncol 19, 3042–3049 (2012).

10. C. H. Lin et al., Microenvironment rigidity modulates responses to the HER2 receptor tyrosine kinase inhibitor lapatinib via YAP and TAZ transcription factors. Mol Biol Cell 26, 3946–3953 (2015).

11. J. P. Fatherree, J. R. Guarin, R. A. McGinn, S. P. Naber, M. J. Oudin, Chemotherapy-Induced Collagen IV Drives Cancer Cell Motility through Activation of Src and Focal Adhesion Kinase. Cancer Res 82, 2031–2044 (2022).

12. J. Cooper, F. G. Giancotti, Integrin Signaling in Cancer: Mechanotransduction, Stemness, Epithelial Plasticity, and Therapeutic Resistance. Cancer Cell 35, 347–367 (2019).

13. G. Halder, S. Dupont, S. Piccolo, Transduction of mechanical and cytoskeletal cues by YAP and TAZ. Nat Rev Mol Cell Biol 13, 591–600 (2012).

14. Z. Meng et al., RAP2 mediates mechanoresponses of the Hippo pathway. Nature 560, 655–660 (2018).

15. S. Dupont et al., Role of YAP/TAZ in mechanotransduction. Nature 474, 179–183 (2011).

16. K. F. Harvey, C. M. Pfleger, I. K. Hariharan, The Drosophila Mst ortholog, hippo, restricts growth and cell proliferation and promotes apoptosis. Cell 114, 457–467 (2003).

17. J. Huang, S. Wu, J. Barrera, K. Matthews, D. Pan, The Hippo signaling pathway coordinately regulates cell proliferation and apoptosis by inactivating Yorkie, the Drosophila Homolog of YAP. Cell 122, 421–434 (2005).

18. S. Wu, J. Huang, J. Dong, D. Pan, hippo encodes a Ste-20 family protein kinase that restricts cell proliferation and promotes apoptosis in conjunction with salvador and warts. Cell 114, 445–456 (2003).

19. F. X. Yu, B. Zhao, K. L. Guan, Hippo Pathway in Organ Size Control, Tissue Homeostasis, and Cancer. Cell 163, 811–828 (2015).

20. Q. Zeng, W. Hong, The emerging role of the hippo pathway in cell contact inhibition, organ size control, and cancer development in mammals. Cancer Cell 13, 188–192 (2008).

21. B. Zhao, L. Li, Q. Lei, K. L. Guan, The Hippo-YAP pathway in organ size control and tumorigenesis: an updated version. Genes Dev 24, 862–874 (2010).

22. Y. Zheng, D. Pan, The Hippo Signaling Pathway in Development and Disease. Dev Cell 50, 264–282 (2019).

23. Q. Chen et al., A temporal requirement for Hippo signaling in mammary gland differentiation, growth, and tumorigenesis. Genes Dev 28, 432–437 (2014).

24. M. Aragona et al., A mechanical checkpoint controls multicellular growth through YAP/TAZ regulation by actin-processing factors. Cell 154, 1047–1059 (2013).

25. T. A. Marlowe, F. L. Lenzo, S. A. Figel, A. T. Grapes, W. G. Cance, Oncogenic Receptor Tyrosine Kinases Directly Phosphorylate Focal Adhesion Kinase (FAK) as a Resistance Mechanism to FAK-Kinase Inhibitors. Mol Cancer Ther 15, 3028–3039 (2016).

26. X. Feng et al., A Platform of Synthetic Lethal Gene Interaction Networks Reveals that the GNAQ Uveal Melanoma Oncogene Controls the Hippo Pathway through FAK. Cancer Cell 35, 457–472 e455 (2019).

27. D. J. Konieczkowski, C. M. Johannessen, L. A. Garraway, A Convergence-Based Framework for Cancer Drug Resistance. Cancer Cell 33, 801–815 (2018).

28. Y. L. Tai et al., An EGFR/Src-dependent beta4 integrin/FAK complex contributes to malignancy of breast cancer. Sci Rep 5, 16408 (2015).

29. L. Zhang et al., EZH2 engages TGFbeta signaling to promote breast cancer bone metastasis via integrin beta1-FAK activation. Nat Commun 13, 2543 (2022).

30. O. Saatci et al., Targeting lysyl oxidase (LOX) overcomes chemotherapy resistance in triple negative breast cancer. Nat Commun 11, 2416 (2020).

31. N. Furth, Y. Aylon, The LATS1 and LATS2 tumor suppressors: beyond the Hippo pathway. Cell Death Differ 24, 1488–1501 (2017).

32. T. Ando et al., EGFR Regulates the Hippo pathway by promoting the tyrosine phosphorylation of MOB1. Commun Biol 4, 1237 (2021).

33. X. H. Yang et al., Disruption of laminin-integrin-CD151-focal adhesion kinase axis sensitizes breast cancer cells to ErbB2 antagonists. Cancer Res 70, 2256–2263 (2010).

34. R. S. Finn et al., Dasatinib, an orally active small molecule inhibitor of both the src and abl kinases, selectively inhibits growth of basal-type/”triple-negative” breast cancer cell lines growing in vitro. Breast Cancer Res Treat 105, 319–326 (2007).

35. F. Huang et al., Identification of candidate molecular markers predicting sensitivity in solid tumors to dasatinib: rationale for patient selection. Cancer Res 67, 2226–2238 (2007).

36. D. Tryfonopoulos et al., Src: a potential target for the treatment of triple-negative breast cancer. Ann Oncol 22, 2234–2240 (2011).

37. W. Guo et al., Beta 4 integrin amplifies ErbB2 signaling to promote mammary tumorigenesis. Cell 126, 489–502 (2006).

38. H. Gao et al., The BMP inhibitor Coco reactivates breast cancer cells at lung metastatic sites. Cell 150, 764–779 (2012).

39. H. Gao et al., Multi-organ Site Metastatic Reactivation Mediated by Non-canonical Discoidin Domain Receptor 1 Signaling. Cell 166, 47–62 (2016).

40. R. Pelossof et al., Prediction of potent shRNAs with a sequential classification algorithm. Nat Biotechnol 35, 350–353 (2017).

41. C. Fellmann et al., An optimized microRNA backbone for effective single-copy RNAi. Cell Rep 5, 1704–1713 (2013).

42. N. E. Sanjana, O. Shalem, F. Zhang, Improved vectors and genome-wide libraries for CRISPR screening. Nat Methods 11, 783–784 (2014).

43. O. Shalem et al., Genome-scale CRISPR-Cas9 knockout screening in human cells. Science 343, 84–87 (2014).

44. K. Labun, M. Krause, Y. Torres Cleuren, E. Valen, CRISPR Genome Editing Made Easy Through the CHOPCHOP Website. Curr Protoc 1, e46 (2021).

45. G. Sebastiaan Winkler et al., Isolation and mass spectrometry of transcription factor complexes. Methods 26, 260–269 (2002).

46. B. Langmead, S. L. Salzberg, Fast gapped-read alignment with Bowtie 2. Nat Methods 9, 357–359 (2012).

47. M. I. Love, W. Huber, S. Anders, Moderated estimation of fold change and dispersion for RNA-seq data with DESeq2. Genome Biol 15, 550 (2014).

48. A. Subramanian et al., Gene set enrichment analysis: a knowledge-based approach for interpreting genome-wide expression profiles. Proc Natl Acad Sci U S A 102, 15545–15550 (2005).

