## Supporting Information for "FAK-SFK Signaling Integrates ECM Rigidity Sensing and Engagement of ERBB2 to Activate YAP and Promote Invasive Growth and Metastasis in Breast Cancer"

This file includes:

Figures S1-S7
Table S1

**Figure Legend:**

**Figure S1.** Correlation analysis between YAP1 expression and prognosis in breast cancer patients. Related to Figure 1.
Kaplan-Meier plot showing YAP1 expression level isn't correlated with poor prognosis in Luminal A, Luminal B and Basal type breast cancer patients.

**Figure S2.** Src induces tyrosine phosphorylation of LATS1. Related to Figure 2.
(A) 293T cells were transfected with FH-LATS1 together with FAK or Src-Y527F. FH-LATS1 was immunoprecipitated and immunoblotted with indicated antibodies.
(B) 293T cells were transfected with FLAG-YAP with FAK or Src-Y527F. FLAG-YAP was immunoprecipitated and immunoblotted with indicated antibodies.
(C) FH-LATS1 was immunoprecipitated from 293T cells after adding different doses of purified GST-Src protein, followed by immunoblotting with indicated antibodies.
(D) HA-LATS1/2 was co-expressed with Src family kinase members, Src, Yes, and Fyn, in 293T cells and immunoprecipitated followed by immunoblotting with indicated antibodies.

**Figure S3.** Src induces LATS1 phosphorylation at Y916/Y1026/Y1076. Related to Figure 3.
(A) Dot blotting of the phosphorylation site specific antibodies targeting p-LATS1-Y916, p-LATS1-Y1026, and p-LATS1-Y1076.
(B) Immunofluorescence imaging of HA-LATS1 (RFP) and YAP (GFP) in LATS1/2 knockout 293A cells re-introduced with LATS1 (single point mutant forms, Y916E, Y1026E, or Y1076E). Scale Bars, 20 $\mu$m.
(C) Immunofluorescence quantification of YAP localization in LATS1/2 knockout 293A cells re-introduced with LATS1 (wild type and different mutant forms). Error bars denote Mean + SD.

(D) Immunofluorescence imaging of HA-LATS1 (RFP) and YAP (GFP) and quantification of YAP localization in Lats1/2 knockout 293A cells re-introduced with wild type LATS1 or mutant LATS1-3YE (Y916/1026/2076E). Scale Bars, 20 μm.

**Figure S4.** Src induces tyrosine phosphorylation of MOB1. Related to Figure 3.
(A) FLAG-MST1/2, HA-SAV were co-transfected with Src or FAK in 293T cells. Immunoprecipitation was performed followed by immunoblotting with indicated antibodies.
(B) Src was transfected with different mutant forms of MOB1 in 293T cells. Immunoprecipitation of HA-MOB1 was performed followed by immunoblotting with indicated antibodies.
(C) All tyrosine sites of MOB1 were mutated to phenylalanine (8YF) and mutated back to tyrosine one by one. Immunoprecipitation of HA-MOB1 was performed followed by immunoblotting with indicated antibodies.
(D) Src, FAK and siFAK were transfected in 293T cells as indicated. Immunoprecipitation was performed followed by immunoblotting with indicated antibodies.

**Figure S5.** YAP activation is required for Src-induced transformation. Related to Figure 5.
(A) GSEA analysis of RNA-seq data in DMSO and Dasatinib treated B77 cells.
(B) MTT assays in Yap knockdown B77 cells.

**Figure S6.** Depletion of Yap/Taz and Fak inhibit tumor growth and invasion in vitro and in vivo. Related to Figure 6.
(A) Immunoblotting of indicated antibodies in inducible Yap/Taz or Ptk2 knockdown MMTV-Neu cells.
(B) Quantification of tumor volumes of female nude mice injected in mammary fat pad with inducible Ptk2 knockdown MMTV-Neu cells. Doxycycline induction began at Week 2 after cell injection. Error bars denote Mean + SD; P values (two-tailed t-test): $^{**}p<0.01$.

(C) Representative Images and normalized photon flux of nude mice injected through tail vein with inducible Ptk2 knockdown MMTV-Neu cells. Doxycycline induction began at Week 1 after cell injection. Error bars denote Mean + SD; P values (two-tailed t-test): *p<0.05.
(D) KEGG analysis of RNA-seq data of lung metastatic tumors from nude mice injected with inducible Ptk2 knockdown MMTV-Neu cells.

**Figure S7.** Dasatinib inhibits HER2 breast cancer growth and metastasis. Related to Figure 7.
(A) Representative imaging of H&E staining and IHC staining of Ki67 and C-Caspase3 in the tumor samples harvested from MMTV-Neu mice after treatment. Scale Bars, 50 μm.
(B) Quantification of IHC staining of Ki67 and C-Caspase3 in the tumor samples harvested from MMTV-Neu mice after treatment. Error bars denote Mean + SD; P values (two-tailed t-test): *p<0.05, **p<0.01.
(C) Representative of IHC staining imaging of Yap and p-Yap (S127) in the tumor samples harvested from MMTV-Neu mice after treatment. Scale Bars, 50 μm.
(D) Quantification of IHC staining of Ki67 and C-Caspase3 in the lung metastases samples harvested from MMTV-Neu cell-injected nude mice after the treatment indicated in Fig. 7E.

**Table S1.** Mass spectrometry mapping results indicating the potential phosphorylation sites on LATS1.

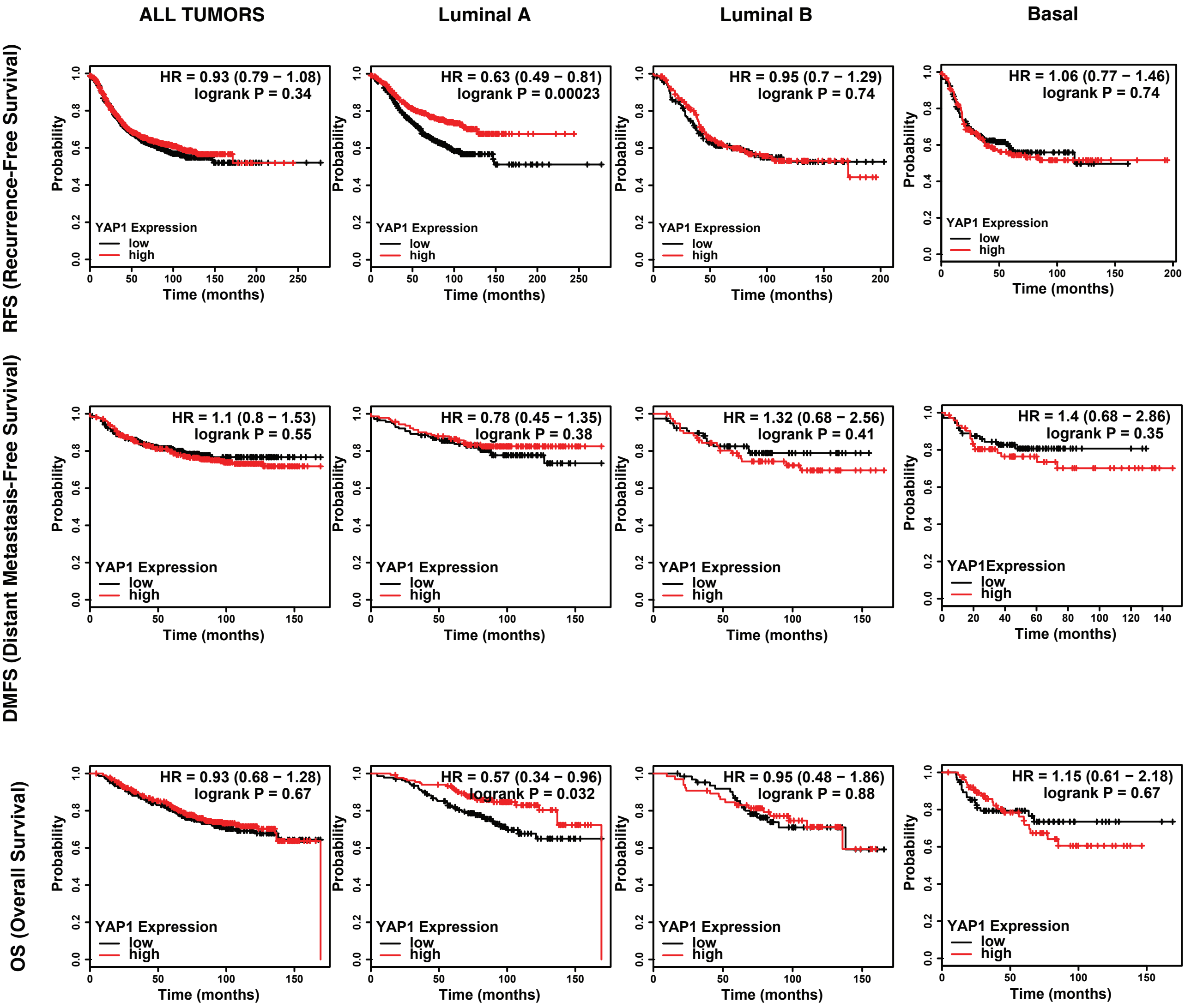
ALL TUMORS
Luminal A
Luminal B
Basal
RFS (Recurrence-Free Survival)
DMFS (Distant Metastasis-Free Survival)
OS (Overall Survival)
HR = 0.93 (0.79 − 1.08)
logrank P = 0.34
HR = 0.63 (0.49 − 0.81)
logrank P = 0.00023
HR = 0.95 (0.7 − 1.29)
logrank P = 0.74
HR = 1.06 (0.77 − 1.46)
logrank P = 0.74
HR = 1.1 (0.8 − 1.53)
logrank P = 0.55
HR = 0.78 (0.45 − 1.35)
logrank P = 0.38
HR = 1.32 (0.68 − 2.56)
logrank P = 0.41
HR = 1.4 (0.68 − 2.86)
logrank P = 0.35
HR = 0.93 (0.68 − 1.28)
logrank P = 0.67
HR = 0.57 (0.34 − 0.96)
logrank P = 0.032
HR = 0.95 (0.48 − 1.86)
logrank P = 0.88
HR = 1.15 (0.61 − 2.18)
logrank P = 0.67
YAP1 Expression
low
high
Probability
Time (months)

Figure S1

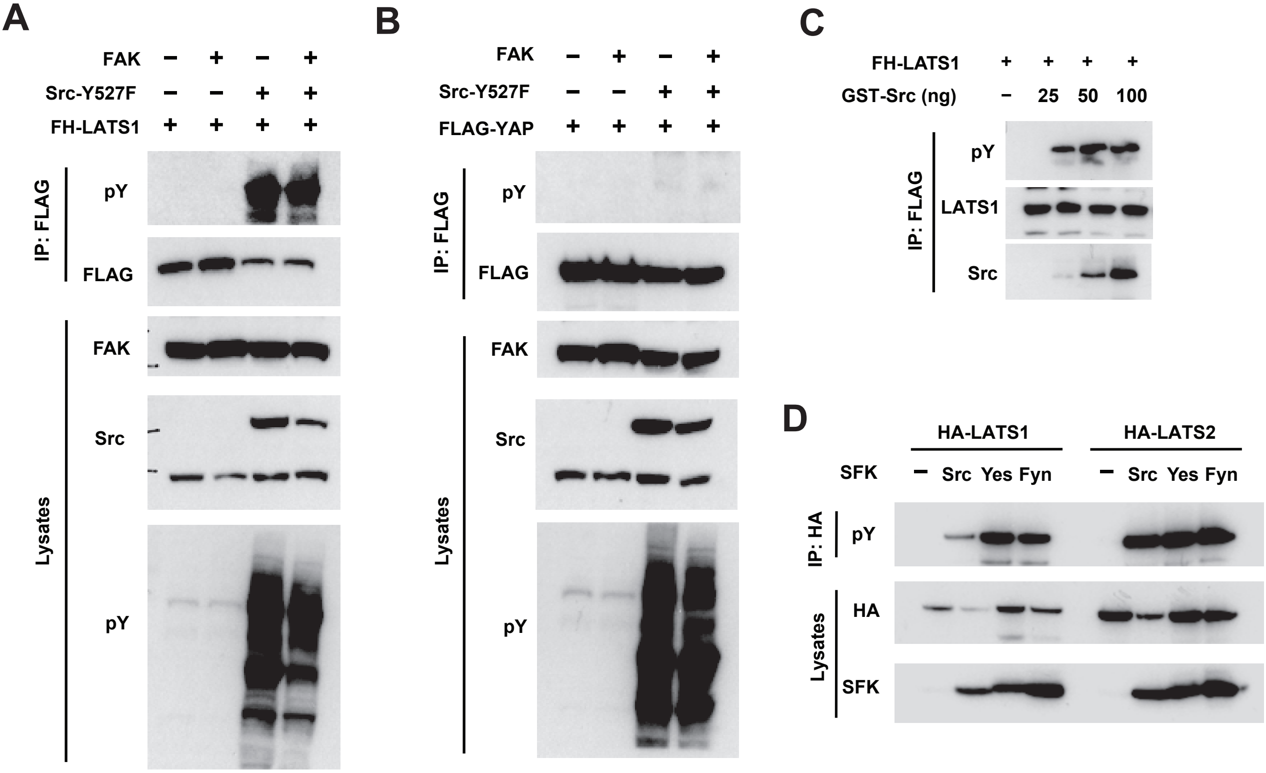
A
FAK − + − +
Src-Y527F − − + +
FH-LATS1 + + + +
IP: FLAG
pY
FLAG
Lysates
FAK
Src
pY
B
FAK − + − +
Src-Y527F − − + +
FLAG-YAP + + + +
IP: FLAG
pY
FLAG
Lysates
FAK
Src
pY
C
FH-LATS1 + + + +
GST-Src (ng) − 25 50 100
IP: FLAG
pY
LATS1
Src
D
HA-LATS1
HA-LATS2
SFK − Src Yes Fyn − Src Yes Fyn
IP: HA
pY
Lysates
HA
SFK

Figure S2

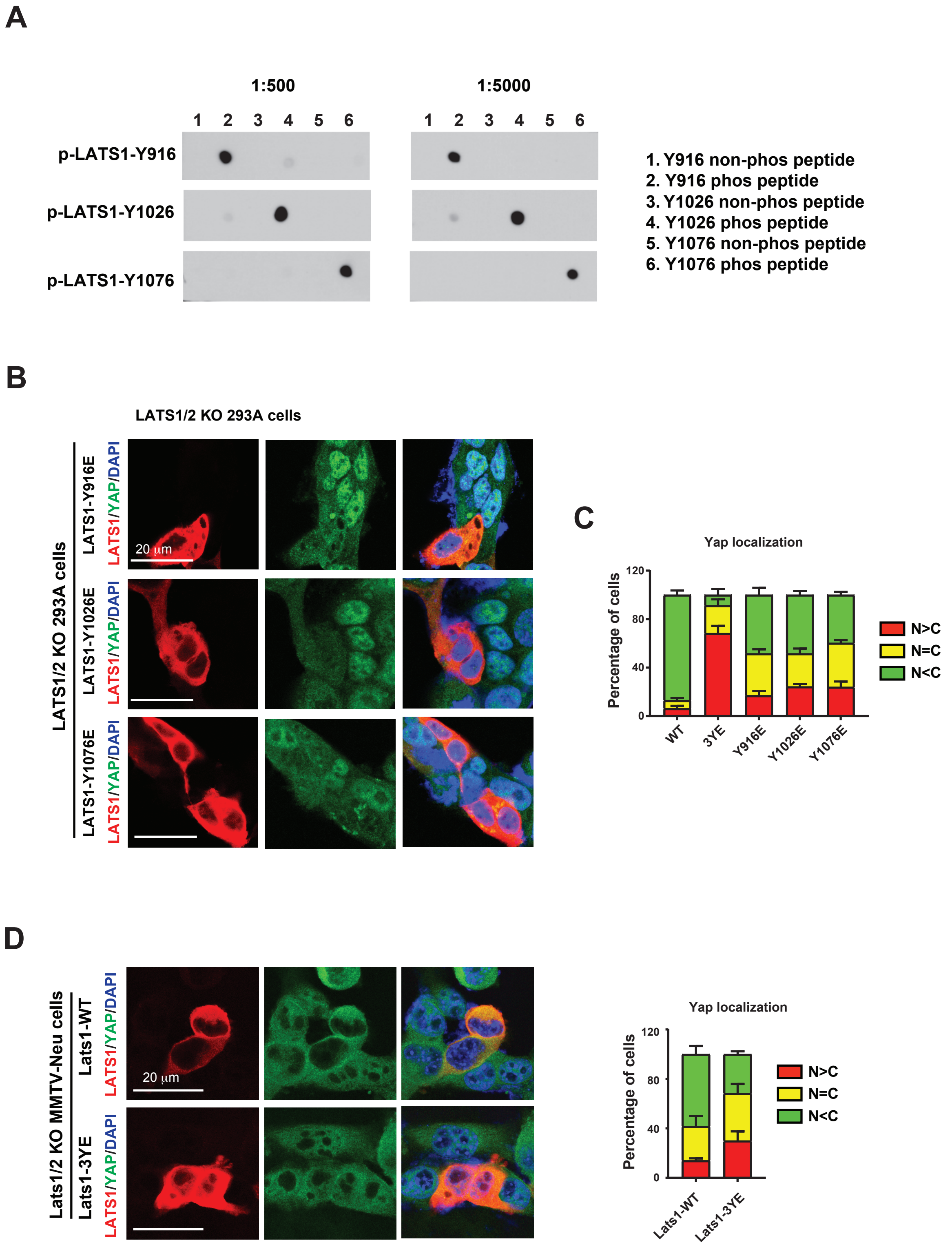
A
1:500
1:5000
1 2 3 4 5 6
p-LATS1-Y916
p-LATS1-Y1026
p-LATS1-Y1076
1. Y916 non-phos peptide
2. Y916 phos peptide
3. Y1026 non-phos peptide
4. Y1026 phos peptide
5. Y1076 non-phos peptide
6. Y1076 phos peptide
B
LATS1/2 KO 293A cells
LATS1/2 KO 293A cells
LATS1-Y916E
LATS1-Y1026E
LATS1-Y1076E
LATS1/YAP/DAPI
20 μm
C
Yap localization
Percentage of cells
120
80
40
0
WT
3YE
Y916E
Y1026E
Y1076E
N>C
N=C
N<C
D
Lats1/2 KO MMTV-Neu cells
Lats1-WT
Lats1-3YE
LATS1/YAP/DAPI
20 μm
Yap localization
Percentage of cells
Lats1-WT
Lats1-3YE
N>C
N=C
N<C

Figure S3

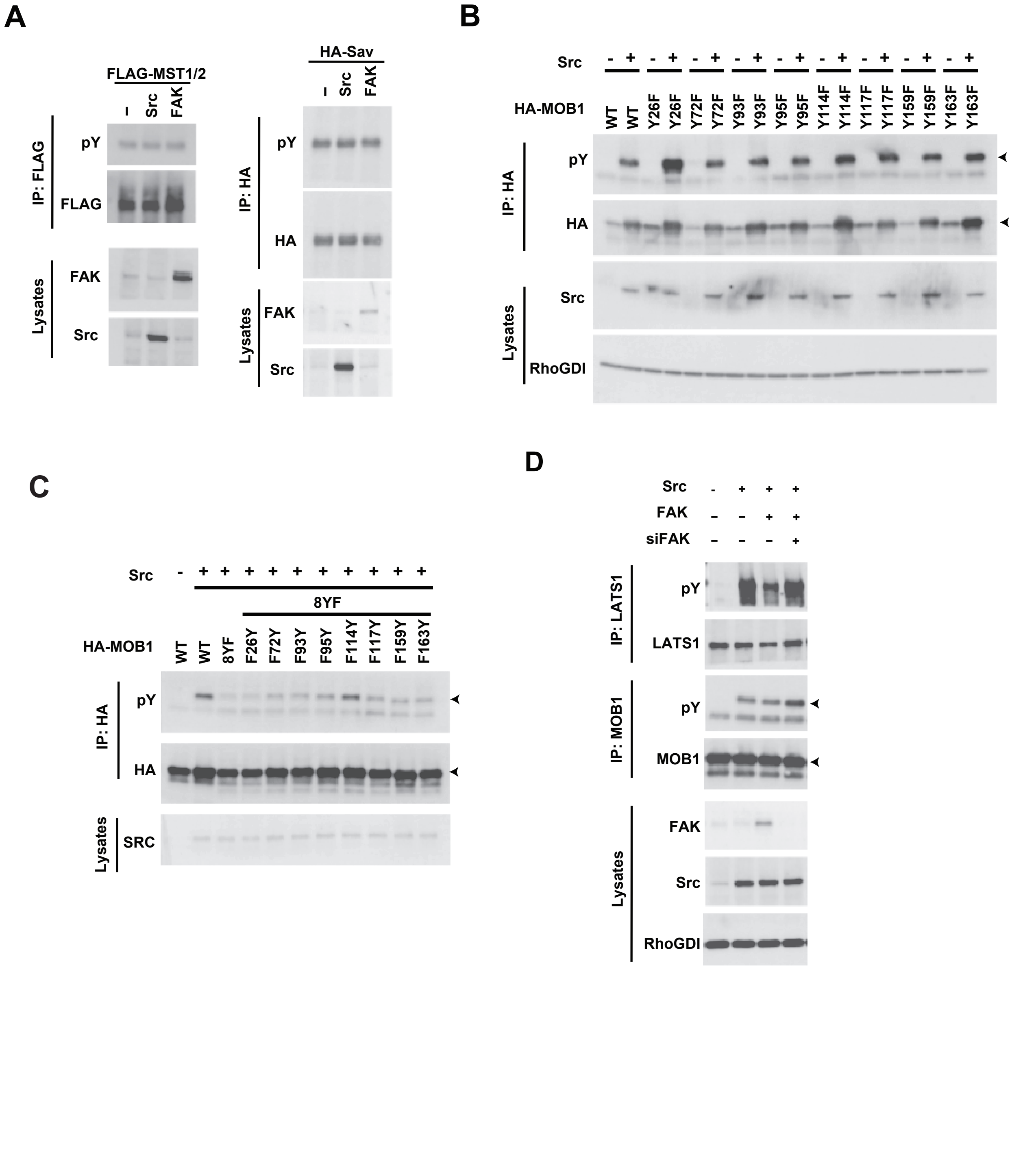
A
FLAG-MST1/2
-
Src
FAK
IP: FLAG
pY
FLAG
Lysates
FAK
Src
HA-Sav
-
Src
FAK
IP: HA
pY
HA
Lysates
FAK
Src
B
Src
- + - + - + - + - + - + - + - + - +
HA-MOB1
WT WT Y26F Y26F Y72F Y72F Y93F Y93F Y95F Y95F Y114F Y114F Y117F Y117F Y159F Y159F Y163F Y163F
IP: HA
pY
HA
Lysates
Src
RhoGDI
C
Src
- + + + + + + + + + +
8YF
HA-MOB1
WT WT 8YF F26Y F72Y F93Y F95Y F114Y F117Y F159Y F163Y
IP: HA
pY
HA
Lysates
SRC
D
Src
- + + +
FAK
- - + +
siFAK
- - - +
IP: LATS1
pY
LATS1
IP: MOB1
pY
MOB1
Lysates
FAK
Src
RhoGDI

Figure S4

**A**

**B77 cells: DMSO vs. Dasatinib**

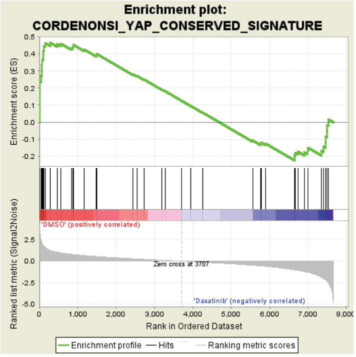


**B**

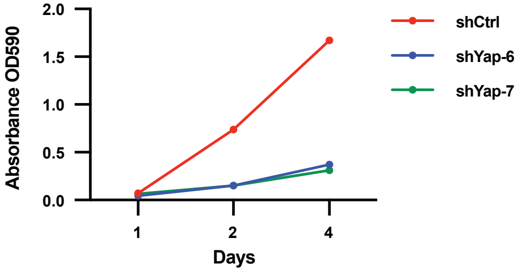


**Figure S5**

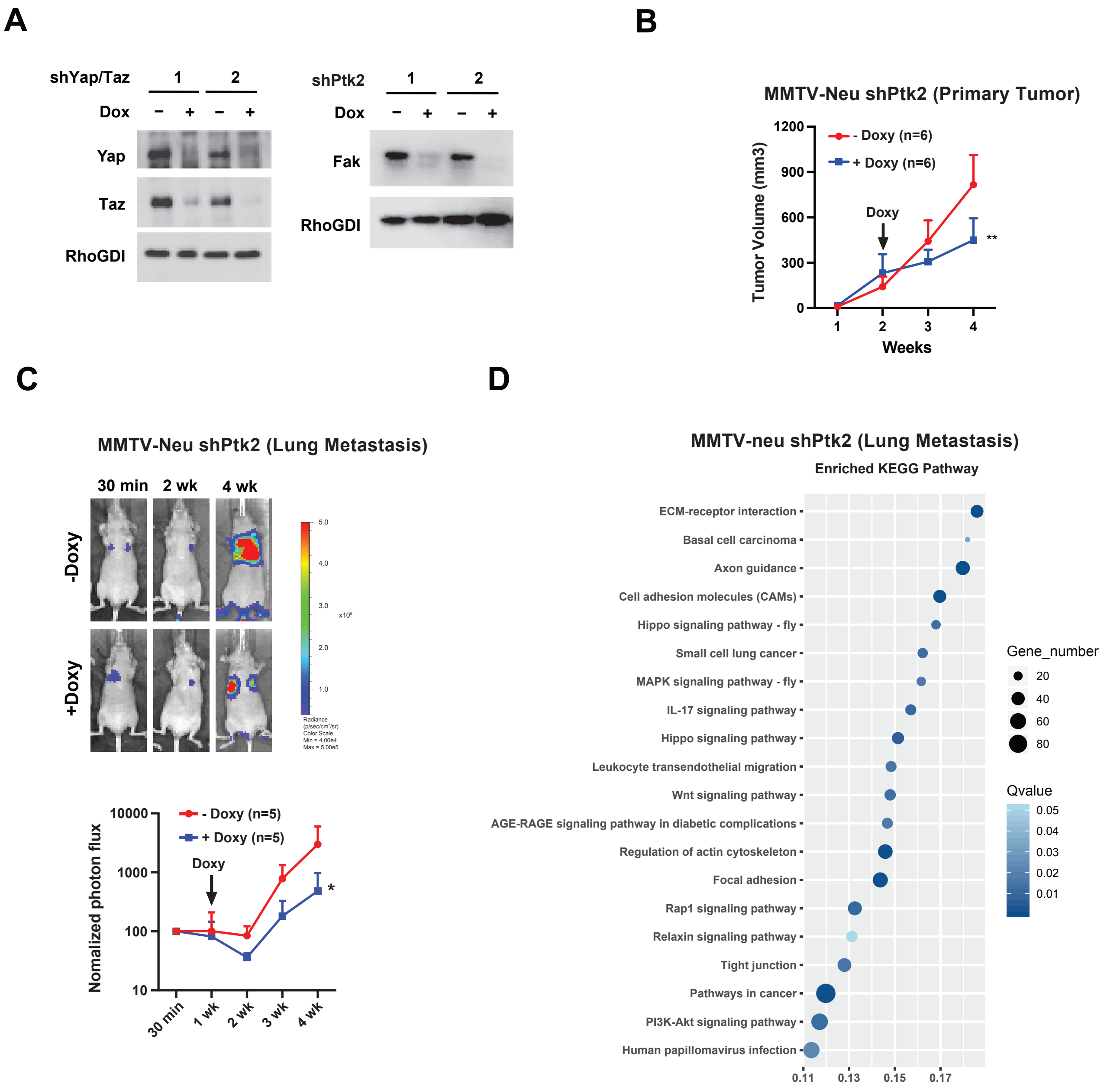
A
shYap/Taz
1
2
Dox
−
+
−
+
Yap
Taz
RhoGDI
shPtk2
1
2
Dox
−
+
−
+
Fak
RhoGDI
B
MMTV-Neu shPtk2 (Primary Tumor)
- Doxy (n=6)
+ Doxy (n=6)
Tumor Volume (mm3)
1200
900
600
300
0
Doxy
**
1
2
3
4
Weeks
C
MMTV-Neu shPtk2 (Lung Metastasis)
30 min
2 wk
4 wk
-Doxy
+Doxy
5.0
4.0
3.0
x10⁵
2.0
1.0
Radiance (p/sec/cm²/sr)
Color Scale
Min = 4.00e4
Max = 5.00e5
- Doxy (n=5)
+ Doxy (n=5)
Nomalized photon flux
10000
1000
100
10
Doxy
*
30 min
1 wk
2 wk
3 wk
4 wk
D
MMTV-neu shPtk2 (Lung Metastasis)
Enriched KEGG Pathway
ECM-receptor interaction
Basal cell carcinoma
Axon guidance
Cell adhesion molecules (CAMs)
Hippo signaling pathway - fly
Small cell lung cancer
MAPK signaling pathway - fly
IL-17 signaling pathway
Hippo signaling pathway
Leukocyte transendothelial migration
Wnt signaling pathway
AGE-RAGE signaling pathway in diabetic complications
Regulation of actin cytoskeleton
Focal adhesion
Rap1 signaling pathway
Relaxin signaling pathway
Tight junction
Pathways in cancer
PI3K-Akt signaling pathway
Human papillomavirus infection
0.11
0.13
0.15
0.17
Gene_number
20
40
60
80
Qvalue
0.05
0.04
0.03
0.02
0.01

**Figure S6**

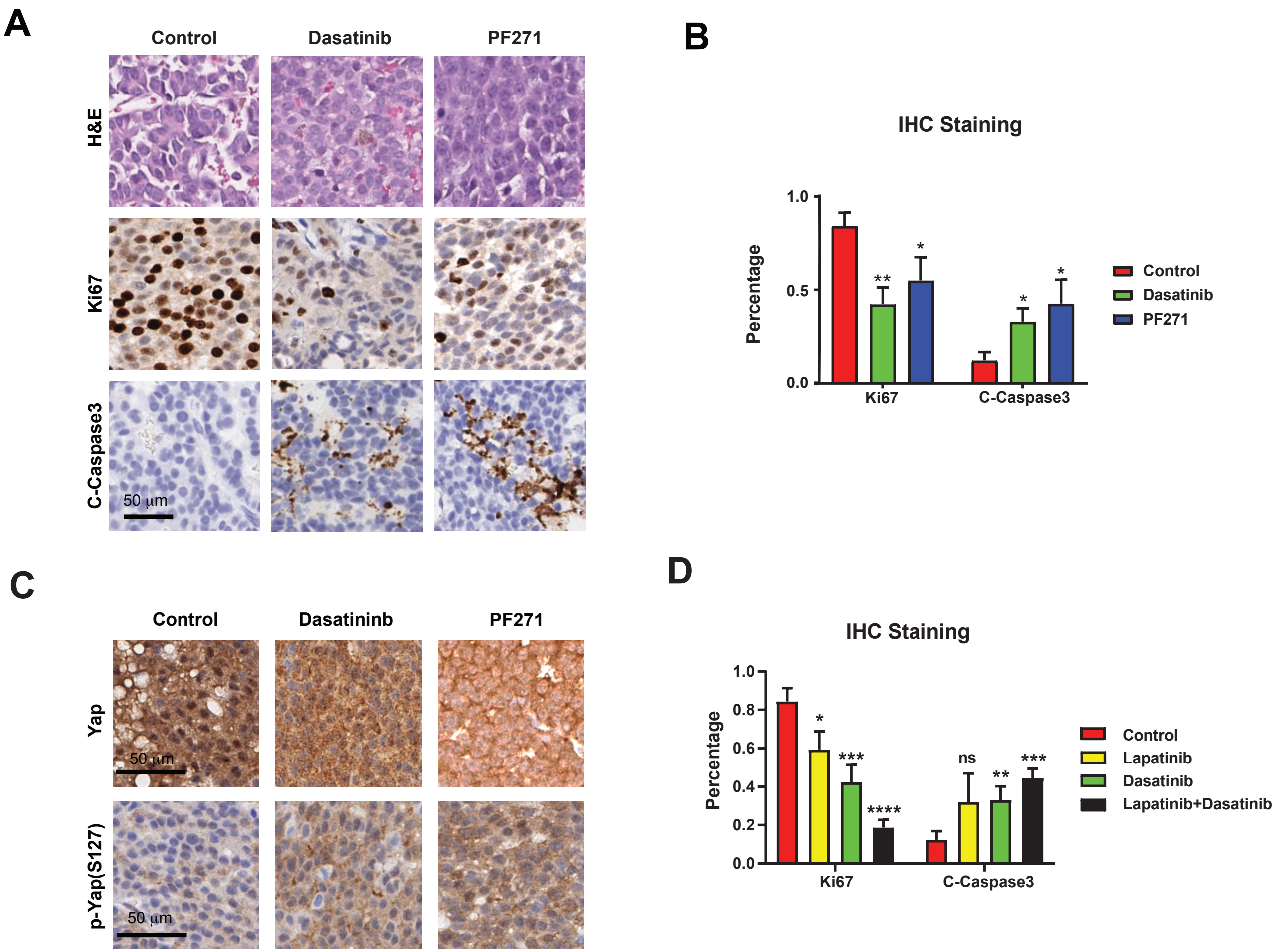
A
Control
Dasatinib
PF271
H&E
Ki67
C-Caspase3
50 µm
B
IHC Staining
Percentage
1.0
0.5
0.0
**
*
*
*
Ki67
C-Caspase3
Control
Dasatinib
PF271
C
Control
Dasatininb
PF271
Yap
p-Yap(S127)
50 µm
50 µm
D
IHC Staining
Percentage
1.0
0.8
0.6
0.4
0.2
0.0
*
***
****
ns
**
***
Ki67
C-Caspase3
Control
Lapatinib
Dasatinib
Lapatinib+Dasatinib

Figure S7

| | | 1st | 2nd |
|---|---|---|---|
| Peptide Sequence: | Residue #: | Ratio of mod/unmod | Ratio of mod/unmod |
| TFPASNYTVSSR | Tyr 23 | 14/136=**0.103** | 7/136=**0.051** |
| HGPPLGESVAYHSESPNSQT | Tyr 200 | 12/66=**0.18** | 1/12=**0.08** |
| RYSGNMEYVISR | Tyr 283 | 36/152=**0.24** | 5/1=**5** |
| RYSGNMEYVISR | Tyr 277 | 7/152=**0.05** | |
| RYSGNMEYVISR | Tyr 277 and 283 | 6/116=**0.05** | |
| EAPNYQGPPPPYPK | Tyr 552 | 1/80=**0.01** | 1/18=**0.056** |
| EAPNYQGPPPPYPK | Tyr 559 | 2/77=**0.03** | |
| HLLHQNPSVPPYESISKPSK | Tyr 573 | 44/168=**0.26** | 21/138=**0.152** |
| EDESEKSYENV | Tyr 597 | 2/13=**0.15** | |
| IQSYSPQAFK | Tyr 634 | 2/116=**0.02** | 2/134=**0.015** |
| LYYSFQDK | Tyr 769 | 1/1=**1** | |
| DKDNLYFVM | Tyr 779 | 26/14=**1.9** | |
| DYIPGGDMMSLLIR | Tyr 784 | 8/13=**0.62** | |
| FYIAELTCAVESVHK | Tyr 808 | 1/82=**0.01** | |
| WTHDSKYYQSG | Tyr861 | 1/1=**1** | |
| CLAHSLVGTPNYIAPEVLLR | Tyr 916 | 1/128=**0.008** | 2/12=**0.17** |
| QQSASYIPK | Tyr 1026 | 1/78=**0.01** | 1/5=**0.2** |
| DTLNGWYK | Tyr 1065 | 7/21=**0.33** | |
| NGKHPEHAFYEFTFR | Tyr 1076 | 10/22=**0.45** | |
| DDNGGYPYNYPKPIEY | Tyr 1092 | 5/13=**0.38** | |
| DDNGGYPYNYPKPIEY | Tyr 1090 | 1/8=**0.13** | |
| NRDLVYV | Tyr 1129 | | 1/2=**0.5** |

**Table S1**
